# Synthesis of stable isotope labeled analogs of phytanic acid for separate and combined tracing of alpha-, beta- and omega-oxidation

**DOI:** 10.1101/2025.11.24.689497

**Authors:** Robert L. Broadrup, Aron C. Schwartz, Julia Smith, Dora von Trentini, Nathaniel W. Snyder

**Affiliations:** Aging + Cardiovascular Discovery Center, Department of Cardiovascular Sciences, Lewis Katz School of Medicine at Temple University; Department of Chemistry, Haverford College

**Keywords:** Phytanic Acid, stable isotope tracing, branched very long chain fatty acid, peroxisome, metabolism Abstract

## Abstract

Phytanic acid (3,7,11,15-tetramethylhexadecanoic acid) is a branched very long chain fatty acid (VLCFA) derived in humans from the diet and catabolized via peroxisomal metabolism. Catabolism of phytanic acid is important in humans as certain inborn errors of peroxisomal metabolism, including Refsum Disease, manifest when catabolism of diet derived phytanic acid is impaired. We developed a novel, linear, unified, unoptimized synthesis of all-racemic phytanic acid along with its 2,3,6,7,10,11,14,15-d8, 1,2-^13^C_2_, and 1,2-^13^C_2_-2,3,6,7,10,11,14,15-d8 stable isotope labeled analogs from commercially available materials including the common starting material ethyl 2-bromoacetate and ethyl 2-bromoacetate-^13^C_2_. This improves upon previous synthetic work to provide a simple synthetic route to produce amounts of stable isotope labeled analogs useful for biochemical studies. The neutron encoded position of the stable isotope enrichment allow distinct examination of oxidative pathways of catabolism, allowing comparison of alpha, beta, and omega oxidation through the combination of labels. This provides a useful set of stable isotope labeled analogs for studies of phytanic acid and branched VLCFA metabolism.

## Introduction

Phytanic acid (PA), 3,7,11,15-tetramethylhexadecanoic acid, is a diet derived metabolic intermediate from the plant derived *trans*-phytol precursor. PA exists as two diastereomers, 3*R*,7*R*,11*R*,15-tetramethylhexadecanoic acid and 3*S*,7*R*,11*R*,15-tetramethylhexadecanoic acid, in a ratio that depends upon their source (1). PA derives from the enzymatic cleavage of phytol from chlorophyll and its subsequent oxidation in the rumen of various animals (2) and in marine organisms. Human metabolism is not know to be capable of degrading chlorophyll to phytol, but studies suggest that phytyl fatty acid esters in vegetables in the human diet may be cleavable in the human digestive system (3). It may then be possible for this phytol to be converted to phytanic acid in the human body by known pathways, the phytanic acid-producing pathway and the dihydrophytol-producing pathway (4). Correspondingly, plasma levels of phytanic acid have been shown to decrease significantly from meat eaters, to lacto-ovo vegetarians and vegans, and finally at their lowest levels in vegans (2). Multiple diets have been identified that help to lower phytanic acid levels, e.g. Westminster diet; however, vitamin screening is recommended by patients on such diets to monitor for any insufficiencies (5).

PA is a highly branched very long chain fatty acid (VLCFA). Due to the multiple methyl branches in PA, catabolism of these phytol derived products requires peroxisomal metabolic pathways for catabolism in mammals. The peroxisome cellular organelle is responsible for the synthesis of bile acids and plasmalogens, glyoxylic acid detoxification, hydrogen peroxide degradation, alpha-oxidation of PA, and beta-oxidation of VLCFAs and fatty acids (6, 7). As a result of this metabolic dependency, impairment of peroxisomal metabolism results in a spectra of inborn errors of metabolism including Refsum Disease. Refsum Disease stems from a single enzyme deficiency caused by a mutation in the gene for phytanoyl-CoA 2-hydroxylase (PHYH), the enzyme responsible for the first intra-peroxisomal step in the alpha-oxidation of PA (8) Refsum can also be caused by minor defects in the peroxisomal biogenesis factor 7 (PEX7) gene, a gene which encodes for a cytosolic receptor for peroxisomal enzymes. (9, 10) Specifically, the PEX7 incorporates the PHYH enzyme and other proteins into peroxisomes. (11, 12) Statistically, a PHYH deficiency is responsible for more than 90% of Refsum cases with the PEX7 deficiency accounting for less than 10%. (13) In individuals without a mutated PHYH gene or minor mutations in PEX7, the methyl branching at the 3-position in phytanic acid prevents it from successfully undergoing beta-oxidation within the peroxisome; consequently, it must first be oxidized from its phytanoyl-CoA form via an alpha-oxidation process in the peroxisome to its 2-hydroxy-3-methyl intermediate. Thereafter it is further oxidized to pristanal and pristanic acid to complete the alpha-oxidation process, and it is then able to move toward the beta-oxidation pathway as some diastereomeric mixture of pristanic acid ((2*R*,6*R*,10*R*)-2,6,10,14-tetramethylpentadecanoic acid and (2*S*,6*R*,10*R*)-2,6,10,14-tetramethylpentadecanoic acid), which notably lacks methyl-branching at the 3-position. In those possessing the PHYH or PEX7 mutations, however, alpha-oxidation is impacted, and high levels of phytanic acid result. (14) An alternate pathway of catabolism exists, primarily in the liver and kidneys, whereby omega-oxidation allows phytanic acid to be oxidized following a multi-step process to 3-methyladipic acid (3-MAA). (14, 15) A wider spectrum of peroxisomal biogenesis disorders, including Zellweger spectrum disorders, caused by mutations in the peroxisomal biogenesis factor genes result in impaired capacity of both the alpha- and omega-oxidation pathways (6, 10, 16).

To help establish the diagnosis, analysis of phytanic acid levels in plasma or serum has traditionally been the first step, followed by enzyme analysis in fibroblasts or molecular genetic testing to confirm the genetic mutation. In cases of Refsum Disease, phytanic acid levels can vary considerably by diet but will often present as elevated for both PHYH and PEX7 mutations; the phytanic acid/pristanic acid ratio will typically present as elevated vs. healthy patients for both mutations. Lastly addition, pipecolic acid concentration will often present as mildly elevated in the case of PHYH mutation. (13) Significantly, phytanic acid levels may also be found in patients suffering from peroxisomal biogenesis disorders or rhizomelic chondrodysplasia punctata type disorder (17). Current treatment options for Refsum Disease focus chiefly on lowering the dietary intake of phytanic acid and the use of plasmapheresis in cases of very high levels of phytanic acid (5).

Chromatographic and mass spectral analyses have extensively been used to determine the presence of indicators of peroxisomal disorders from specimens including urine, plasma, and dried blood spots (18–21). This has included both gas chromatography-mass spectrometry (GC-MS) and liquid chromatography-mass spectrometry (LC-MS) based approaches to measure phytanic acid and other analytes indicative of peroxisomal disorders. Biochemical studies using radioactive tracers of PA are also reported. Hoefler et al used 1-^14^C-phytanic acid in the form of ammonium-^14^C-phytanate to investigate the oxidation of phytanic acid in cultured skin fibroblasts from patients with a number of peroxisomal disorders and detected reduced phytanic acid oxidation and plasmalogen biosynthesis relative to control in the fibroblasts from RCDP, Zellweger Syndrome, and Neonatal Adrenoleukodystrophy patients by trapping and quantifying ^14^CO_2_ trapped on filters (17). Similarly Kase and Björkhem demonstrated that phytanic acid oxidation was severely impaired in fibroblasts from children with peroxisomal disorders by incubating fibroblasts with 1-^14^C- and U-^3^H-labelled phytanic acid and assaying trapped ^14^CO_2_ as well as hydro- and lipophilic fibroblast extracts for radioactivity (22).

Due to the complex metabolism of PA, considerable work has focused on synthesis of PA and its analogs. Unlabeled phytanic acid is most commonly made from phytol, where Schröder et al provide a synthesis of both natural diastereomeric phytanic acid from natural 7*R*,11*R*-*trans*-phytol as well as all-racemic-phytanic acid from racemic phytol. Their synthesis uses a hydrogenation of phytol to dihydrophytol, followed by oxidation of the primary alcohol to the carboxylic acid; this approach is applicable to both the stereoselective and all-racemic versions of the synthesis as it leaves the chiral centers in natural 7*R*,11*R*-*trans*-phytol unaffected (1). Similar syntheses not discussed by Schroder et al are referenced here but will not be discussed in detail (23–25). Lastly, Serebryakov et al report the synthesis of all-racemic phtanic acid by a different method. Their synthesis utilizes a Horner-Wadsworth-Emmons Olefination between all-*rac*-6,10,14-trimethylpentadecan-2-one and triethylphosphonoacetate to obtain 3,7,11,15-tetramethylhexadec-2-enoic acid ester, which is subsequently saponified to the corresponding acid and hydrogenated to yield all-racemic phytanic acid (26).

Previous reports of labeled PA include both stable and radioactive isotopes. The Zenger-Hain et al synthesis of 2,3-^3^H-phytanic acid (27) and Stewart & Kates’ (28) synthesis of 1,1-^2^H_2_-phytanol utilize reduction and oxidation chemistry similar to that of Schröder (1). Muñoz and Guerrero (29) shift the position of di-deuteration in their all-racemic 5,5-^2^H_2_-phytol by making use of an LiAlD_4_ reduction of 3,7,11-trimethyldodecanoic acid with subsequent conversion of the primary alcohol to a bromide, conversion to the organocuprate, and epoxide opening. Yepuri extend upon Schröder et al’s process to generate perdeuterated phytanic acid from a diastereomeric mixture of the naturally occurring phytanic acid diastereomers by use of a hydrothermal deuteration with NaOD in D_2_O with Pt/C at 220 °C (25). Lastly, Poulos & Barone (30) and Fingerhut (31) prepare 1-^14^C-phytanic acid by the basic hydrolysis of pristanyl 14C nitrile formed from the reaction of pristanyl iodide and 14C-cyanide; it is unclear from their procedure whether the corresponding product is racemic. Here we expand upon these previous synthetic approaches to generate all-racemic phytanic acid along with its 2,3,6,7,10,11,14,15-d8, 1,2-13C2, and 1,2-13C2-2,3,6,7,10,11,14,15-d8 analogs.

## Materials

The deuterium (^2^H_2_) used in the hydrogenation reactions was 99.8 atom % D, purchased from Sigma Aldrich. The ethyl bromoacetate (1,2-^13^C_2_ 99%) was purchased from Cambridge Isotope Laboratories. All other solvents and reagents were purchased from Sigma Aldrich unless otherwise specified and used as purchased without further purification.

Reaction mixtures were monitored using thin layer chromatography (TLC) on silica gel glass plates. Compounds were visualized using UV light (254 nm), potassium permanganate, or bromocresol green stain. NMR spectra were obtained on a Varian/Agilent 500 MHz NMR spectrometer. NMR samples were dissolved in CDCl_3_, and TMS was used as the internal standard. Chemical shifts (δ) are reported in parts per million (ppm). IR spectra were obtained using a ThermoScientific Nicolet iS5 FTIR with an iD7 ATR attachment. GC/MS spectra were obtained on a Perkin Elmer GAs Chromatograph Clarus 580. Samples for GC/MS were prepared in chloroform or methanol. Detailed spectral characterization of all compounds and intermediates is provided in the supporting information.

## Methods

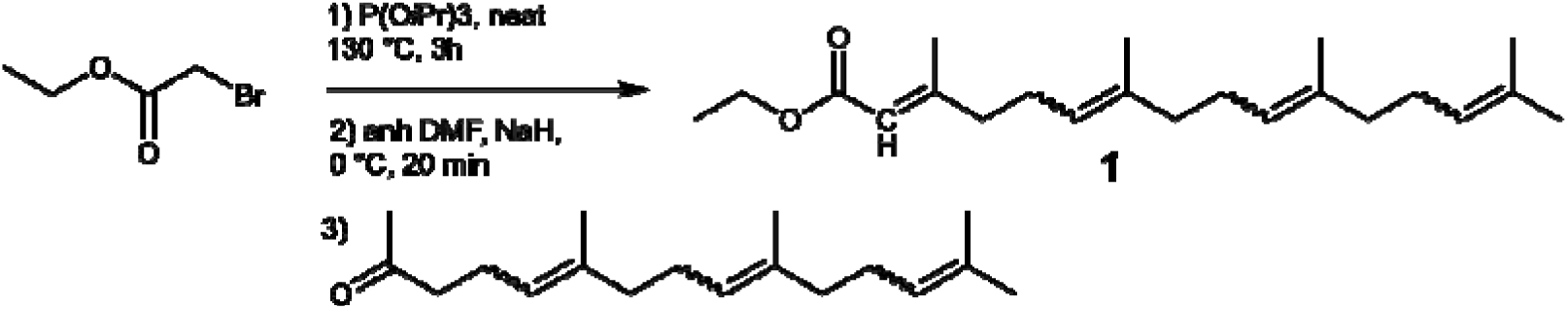

### ethyl (2E)-3,7,11,15-tetramethylhexadeca-2,6,10,14-tetraenoate (1)

The light in the fume hood was turned off for this reaction until it was quenched. To a 100 mL 24/40 single necked round bottom flask with reflux condenser and rubber septum under argon were added ethyl 2- bromoacetate (3.182 g, 19.05 mmol, 2.50 equiv) and triisopropylphospite (9.40 mL, 38.10 mmol, 5.00 equiv) to give a clear, light golden mixture. The mixture was heated at reflux (approximately 130 °C) for 1.5 hr, and then the temperature was increased to 140-145 °C for an additional 2 hr. The mixture was cooled to approximately 0 °C with ice/water bath, and anhydrous DMF (22 mL) was added to the flask. NaH (0.991 g, 24.77 mmol, 3.25 equiv) was then added portionwise as a 60% dispersion in mineral oil at approximately 0 °C to give a grey slurry, which turned to a translucent brown after stirring for 30 minutes. 6,10,14- trimethylpentadeca-5,9,13-trien-2-one (2.00 g, 7.62 mmol, 1.00 equiv) in anhydrous DMF (8 mL) was added dropwise at 2 drops/second. The mixture was allowed to warm to room temperature with stirring overnight to give a translucent orange solution. To the flask was then added ammonium chloride followed by ethyl acetate. The mixture was extracted with ethyl acetate (3X30 mL). The organic layers were combined, washed 5 times with water, washed with brine, dried over magnesium sulfate, filtered and concentrated by rotary evaporation to yield a yellow oil. The crude product was dissolved in a minimal amount of DCM, adsorbed onto dry silica, volatiles removed and loaded onto a silica column packed in hexanes. The product was eluted with several column lengths of 10% ethyl acetate in hexanes to yield a clear/yellow oil (2.12 g, 6.38 mmol, 84% yield). GC-MS *m/z* = 332.03 [M]^+^. ^1^H NMR (500 MHz, CDCl_3_): δ 5.65 (s, 1H), 5.09 (m, 4H), 4.14 (m, 2H), 2.15 (m, 6H), 2.12 (s, 2H), 2.04 (m, 9H), 1.98, (m, 4H), 1.67, (m, 8H), 1.59 (t, 10H). ^13^C NMR (126 MHz, CDCl_3_): δ 208.63, 166.80, 159.62, 136.35, 134.98, 124.35, 122.89, 115.64, 59.37, 43.73, 40.92, 39.68, 31.96, 26.74, 23.32, 17.62, 14.29.

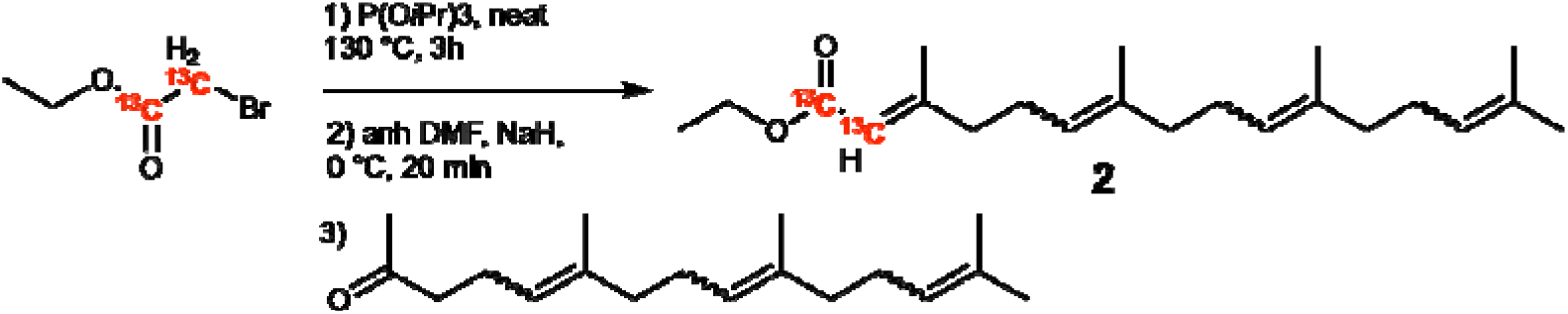

### ethyl (2E)-3,7,11,15-tetramethylhexadeca-2,6,10,14-tetraenoate-1,2-13C2 (2)

The light in the fume hood was turned off for this reaction until it was quenched. To a 100 mL 24/40 single necked round bottom flask with reflux condenser and rubber septum under argon, were added ethyl ethyl 2-bromoacetate-13C2 (3.182 g, 19.05 mmol, 2.50 equiv) and triisopropylphospite (9.40 mL, 38.10 mmol, 5.00 equiv) to give a clear, light golden mixture. The mixture was heated at reflux (approximately 130 °C) for 1.5 hr and then the temperature was increased to 140-145 °C for an additional 2 hr. The mixture was cooled to approximately 0 °C with ice/water bath, and anhydrous DMF (22 mL) was added to the flask. NaH (0.991 g, 24.77 mmol, 3.25 equiv) was then added as a 60% dispersion in mineral oil at approximately 0 °C to give a grey slurry, which turned to a translucent brown after stirring for 30 minutes. 6,10,14- trimethylpentadeca-5,9,13-trien-2-one (2.00 g, 7.62 mmol, 1.00 equiv) in anhydrous DMF (8 mL) was added dropwise at 2 drops/second. The mixture was allowed to warm to room temperature with stirring overnight to give a translucent orange solution. To the flask was added ammonium chloride followed by ethyl acetate. The mixture was extracted with ethyl acetate (3X30 mL). The organic layers were combined, washed 5 times with water, washed with brine, dried over magnesium sulfate, filtered and concentrated to yield a yellow oil. The crude product was dissolved in a minimal a ount of DCM, adsorbed onto dry silica, volatiles removed and dry transferred onto a silica column packed in hexanes. The product was eluted with several column lengths of 10% ethyl acetate in hexanes to yield a clear/yellow oil (2.12 g, 6.38 mmol, 84% yield). GC-MS *m/z* = 334.02 [M]^+^. ^1^H NMR (500 MHz, CDCl_3_): δ 5.81 (s, 1H), 5.49 (s, 1H), 5.09 (t, 4H), 4.12 (m, 2H), 2.65 (m, 1H), 2.44 (m, 1H), 2.26 (q, 1H), 2.15 (s, 7H), 2.04 (m, 7H), 1.97 (m, 4H), 1.67 (s, 7H), 1.59 (s, 8H), 1.24 (m, 3H). ^13^C NMR (126 MHz, CDCl_3_): δ167.10, 136.11, 124.09, 115.94, 59.37, 43.74, 41.85, 39.71, 37.85, 31.96, 26.74, 23.32, 17.62, 14.28.

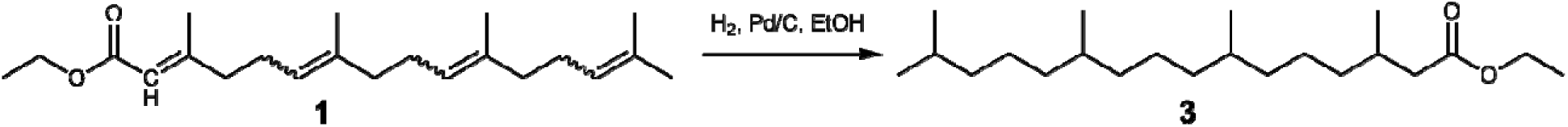

### ethyl 3,7,11,15-tetramethylhexadecanoate (3)

To a round bottom flask were added ethyl (2*E*)-3,7,11,15-tetramethylhexadeca-2,6,10,14-tetraenoate (**1**) (250 mg, 0.752 mmol, 1.00 equiv) dissolved in 200 proof ethanol (3 mL). The round bottom was evacuated and backfilled with argon several times, and then palladium on carbon (10%, 25 mg) was added to the flask. It was evacuated and backfilled with argon 3 times followed by hydrogen gas. The reaction mixture was left stirring overnight, and the balloon was deflated in the morning. The hydrogen balloon was removed, and the flask was evacuated and backfilled with argon 3 times. The reaction mixture was filtered through a pad of celite, and concentrated under reduced pressure. The crude product was dissolved in a minimal amount of DCM, adsorbed onto dry silica, volatiles removed and dry transferred onto a silica column packed in hexanes. The product was eluted with a gradient system starting with 10:90 ethyl acetate:hexanes and increasing to 50:50 ethyl acetate:hexanes to yield a colorless oil (90.2 mg, 35% yield). GC-MS *m/z* = 340.18 [M]^+^. ^1^H NMR (500 MHz, CDCl_3_): δ 4.11 (q, 2H), 2.39 (t, 1H), 2.27 (m, 1H), 2.07 (m, 1H), 1.95 (m, 1H), 1.52 (m, 2H), 1.25 (m, 20H), 0.93 (m, 2H), 0.86 (m, 15H). ^13^C NMR (126MHz, CDCl_3_): δ 209.18, 173.30, 59.99, 44.11, 41.90, 39.35, 37.36, 32.74, 30.39, 27.95, 24.77, 22.67, 21.44, 19.64, 14.25.

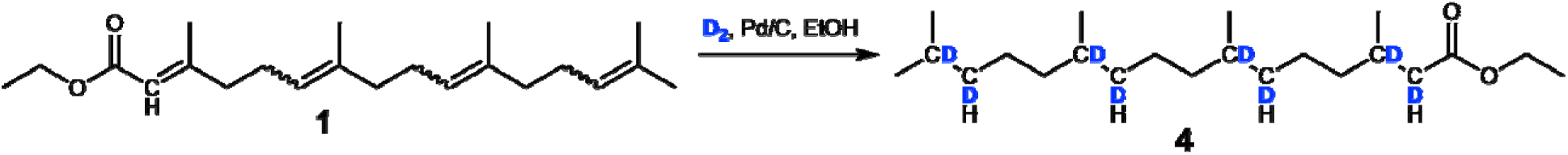

### ethyl 3,7,11,15-tetramethylhexadecanoate-2,3,6,7,10,11,14,15-d8 (4)

To a round bottom flask were added ethyl (2*E*)-3,7,11,15-tetramethylhexadeca-2,6,10,14-tetraenoate (***1***) (250 mg, 0.752 mmol, 1.00 equiv) dissolved in 200 proof ethanol (3 mL). The round bottom was evacuated and backfilled with argon several times and then palladium on carbon (10%, 25 mg) was added to the flask. It was evacuated and backfilled with argon 3 times followed by deuterium gas. The reaction mixture was left stirring overnight, and the balloon was deflated in the morning. The deuterium balloon was removed, and the flask was evacuated and backfilled with argon 3 times. The reaction mixture was filtered through a pad of celite, and concentrated under reduced pressure. The crude product was dissolved in a minimal amount of DCM, adsorbed onto dry silica, volatiles removed and dry transferred onto a silica column packed in hexanes. The product was eluted with a gradient system starting with 10:90 ethyl acetate:hexanes and increasing to 50:50 ethyl acetate:hexanes to yield a colorless oil (186 mg, 0.53 mmol, 71% yield). GC-MS *m/z*= 348.26 [M]^+^. ^1^H NMR (500 MHz, CDCl_3_): δ 4.11 (q, 2H), 2.39 (q, 1H), 2.29 (m, 1H), 2.05 (m, 1H), 1.91 (m, 1H), 1.52 (m, 2H), 1.25 (m, 16H), 0.92 (t, 3H) 0.84 (m, 11H). ^13^C NMR (126 MHz, CDCl_3_): δ 209.09, 173.28, 59.96, 44.09, 41.88, 39.35, 37.04, 29.74, 27.93, 24.28, 22.57, 21.42, 19.61, 14.24.

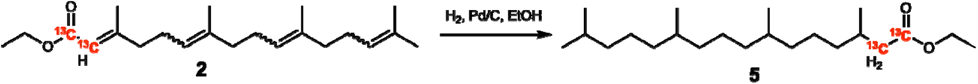

### ethyl 3,7,11,15-tetramethylhexadecanoate-1,2-13C2 (5)

To a round bottom flask were added ethyl (2*E*)-3,7,11,15-tetramethylhexadeca-2,6,10,14-tetraenoate*-1,2-13C2* (**2**) (250 mg, 0.747 mmol, 1.00 equ v) dissolved in 200 proof ethanol (3 mL). The round bottom was evacuated and backfilled with argon several times and then palladium on carbon (10%, 25 mg) was added to the flask. It was evacuated and backfilled with argon 3 times followed by hydrogen gas. The reaction mixture was left stirring overnight, and the balloon was deflated in the morning. The hydrogen balloon was removed, and the flask was evacuated and backfilled with argon 3 times. The reaction mixture was filtered through a pad of celite, and concentrated under reduced pressure. The crude product was dissolved in a minimal amount of DCM, adsorbed onto dry silica, volatiles removed and loaded onto a silica column packed in hexanes. The product was eluted with a gradient system starting with 10:90 ethyl acetate:hexanes and increasing to 50:50 ethyl acetate:hexanes to yield a colorless oil (109 mg, 0.318 mmol, 33% yield). GC-MS *m/z* = 342.16 [M]^+^. ^1^H NMR (500 MHz, CDCl_3_): δ 4.10 (q, 2H), 2.38 (t, 1H), 2.15 (m, 1H), 2.11 (s, 1H), 1.93 (m, 1H), 1.51 (m, 2H), 1.24 (m, 20H), 0.92 (t, 3H), 0.84 (m, 15H). ^13^C NMR (126 MHz, CDCl_3_): δ 173.03, 59.96, 42.49, 39.34, 37.07, 32.75, 27.93, 24.75, 22.57, 19.62, 14.22.

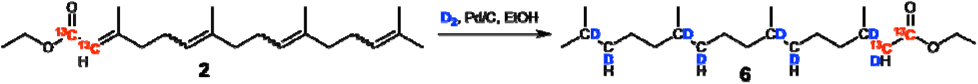

### ethyl 3,7,11,15-tetramethylhexadecanoate-1,2-^13^C_2_-2,3,6,7,10,11,14,15-d_8_ (6)

To a round bottom flask were added ethyl (2*E*)-3,7,11,15-tetramethylhexadeca-2,6,10,14-tetraenoate-*1,2-13C2.* (**2**) (250 mg, 0.747 mmol, 1.00 equiv) dissolved in 200 proof ethanol (3 mL). The round bottom was evacuated and backfilled with argon several times and then palladium on carbon (10%, 25 mg) was added to the flask. It was evacuated and backfilled with argon 3 times followed by deuterium gas. The reaction mixture was left stirring overnight, and the balloon was deflated in the morning. The deuterium balloon was removed, and the flask was evacuated and backfilled with argon 3 times. The reaction mixture was filtered through a pad of celite, and concentrated under reduced pressure. The crude product was dissolved in a minimal amount of DCM, adsorbed onto dry silica, volatiles removed and dry transferred onto a silica column packed in hexanes. The product was eluted with a gradient system starting with 10:90 ethyl acetate:hexanes and increasing to 50:50 ethyl acetate:hexanes to yield a colorless oil (141 mg, 0.402 mmol, 54% yield). GC-MS *m/z* = 351.30 [M]^+^. ^1^H NMR (500 MHz, CDCl_3_): δ 4.12 (m, 2H), 2.39 (t, 1H), 2.20 (m, 1H), 1.95 (m, 1H), 1.51 (m, 1H), 1.25 (m, 17H), 0.93 (t, 3H), 0.86 (m, 13H). ^13^C NMR (126 MHz, CDCl_3_): δ 173.12, 65.86, 60.02, 41.68, 27.96, 22.68, 19.66, 14.27.

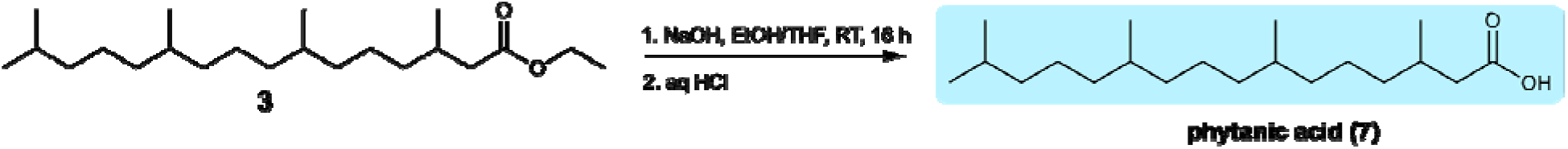

### 3,7,11,15-tetramethylhexadecanoic acid (Phytanic acid). (7)

To a round bottom flask were added ethyl 3,7,11,15-tetramethylhexadecanoate (**3**) (582 mg, 1.71 mmol, 1.00 equiv) dissolved in ethanol (4.35 mL) and THF (4.35 mL) at room temperature to give a clear colorless solution with stirring. NaOH (1.9 mL, 10% aq) was then added quickly to the flask, which was then rubber-stoppered loosely and left to stir for 16 h. To the flask, was then added HCl (2M aqueous, approximately 15 mL) until the pH was 1 by pH strips. DCM was added to the flask, and the mixture was extracted with DCM (3 x 10 mL). The organic layers were combined and dried over MgSO_4_, filtered, and the solvent was removed under reduced pressure to yield a clear oil. The crude product was dissolved in a minimal amount of DCM, adsorbed onto dry silica, volatiles removed and dry transferred onto a silica column packed in hexanes. The product was eluted with 1 column length hexanes, and several column lengths of 10:90 ethyl acetate:hexanes. The solvent was removed under reduced pressure to yield a clear oil (0.25 g, 0.74 mmol, 47% yield). ^1^H NMR (500 MHz, CDCl_3_): δ 2.35 (m,1H), 2.12 (m, 1H), 1.95 (m, 1H), 1.52 (m, 1H), 1.26 (m, 20H), 0.96 (d, 3H), 0.87, (q, 13H). ^13^C NMR (126 MHz, CDCl_3_): δ 65.86, 41.38, 39.36, 37.27, 32.76, 30.18, 27.97, 24.32, 22.62, 19.64. Note: this procedure and data is from Dora von Trentini, using a different starting material batch.

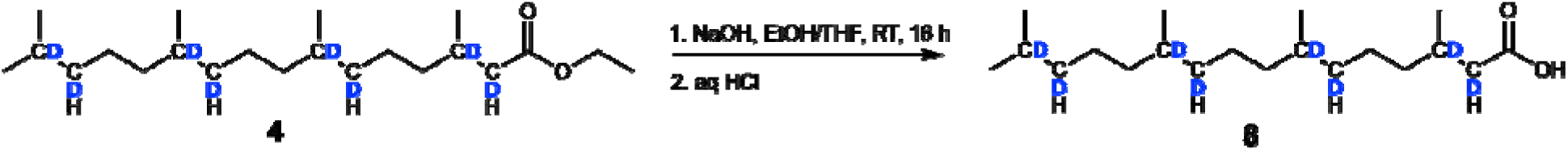

### 3,7,11,15-tetramethylhexadecanoic-2,3,6,7,10,11,14,15-d8 acid (8)

To a round bottom flask were added ethyl 3,7,11,15-tetramethylhexadecanoate-d8 (**4**) (186 mg, 0.530 mmol, 1.00 equiv) dissolved in ethanol (1.36 mL) and THF (1.36 mL) at room temperature to give a clear colorless solution with stirring. NaOH (0.59 mL, 10% aq) was then added quickly to the flask, which was then rubber-stoppered loosely and left to stir for 16 h. To the flask, was then added HCl (2M aqueous, approximately 1.5 mL) until the pH was 1 by pH strips. DCM (4 mL) was added to the flask, and the mixture was extracted with DCM (3 x 4 mL). The organic layers were combined and dried over MgSO_4_, filtered, and the solvent was removed under reduced pressure to yield a clear oil. The crude product was dissolved in a minimal amount of DCM, adsorbed onto dry silica, volatiles removed and dry transferred onto a silica column packed in hexanes. The product was eluted with 1 column length hexanes, and 3 column lengths of 10:90 ethyl acetate: hexanes. The solvent was removed under reduced pressure to yield a clear oil (16.8 mg, 0.052 mmol, 10% yield). GC-MS *m/z* = 319.35 [M]^+^. ^1^H NMR (500 MHz, CDCl_3_): δ 2.33 (dd, 1H), 2.15 (m, 1H), 1.95 (m, 1H), 1.51 (m, 1H), 1.26 (m, 16H), 0.86 (m, 12 H). ^13^C NMR (126 MHz, CDCl_3_): δ 41.29, 37.02, 31.90, 29.33, 24.78, 22.68, 19.51, 14.07. *3,7,11,15-tetramethylhexadecanoic-1,2-13C2 acid* (***9***). To a round bottom flask were added ethyl 3,7,11,15-tetramethylhexadecanoate-1,2-^13^C_2_-2,3,6,7,10,11,14,15-d_8_ (**5**) (141 mg, 0.402 mmol, 1.00 equiv) dissolved in ethanol (1.00 mL) and THF (1.00 mL) at room temperature to give a clear colorless solution with stirring. NaOH (0.45 mL, 10% aq) was then added quickly to the flask, which was then rubber-stoppered loosely and left to stir for 16 h. To the flask, was then added HCl (2M aqueous, approximately 1 mL) until the pH was 1 by pH strips. DCM (4 mL) was added to the flask, and the mixture was extracted with DCM (3 x 3 mL). The organic layers were combined and dried over MgSO_4_, filtered, and the solvent was removed under reduced pressure to yield a clear oil. The crude product was dissolved in a minimal amount of DCM, adsorbed onto dry silica, volatiles removed and dry transferred onto a silica column packed in hexanes. The product was eluted with 3 column lengths of 10:90 ethyl acetate: hexanes. The solvent was removed under reduced pressure to yield a clear oil (61.2 mg, 0.19 mmol, 47% yield). GC-MS *m/z* = 314.31 [M]^+^. ^1^H NMR (500 MHz, CDCl_3_): δ 2.48 (m, 1H), 2.25 (m, 1H), 2.00, (m, 1H), 1.53 (m, 1H), 1.26 (m, 21H), 0.86 (m, 16H). ^13^C NMR (126 MHz, CDCl_3_): δ 177.90, 65.86, 4108, 39.36, 37.37, 32.78, 31.90, 29.33, 27.96, 24.77, 22.68, 19.64, 14.07.

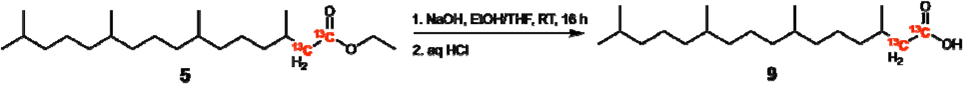

### 3,7,11,15-tetramethylhexadecanoic-1,2-13C2-2,3,6,7,10,11,14,15-d8 acid (10)

To a round bottom flask were added ethyl 3,7,11,15-tetramethylhexadecanoate-1,2-13C2 (**6**) (109 mg, 0.318 mmol, 1.00 equiv) dissolved in ethanol (0.82 mL) and THF (0.82 mL) at room temperature to give a clear colorless solution with stirring. NaOH (0.35 mL, 10% aq) was then added quickly to the flask, which was then rubber-stoppered loosely and left to stir for 16 h. To the flask, was then added HCl (2M aqueous, approximately 1 mL) until the pH was 1 by pH strips. DCM (4 mL) was added to the flask, and the mixture was extracted with DCM (3 x 3 mL). The organic layers were combined and dried over MgSO_4_, filtered, and the solvent was removed under reduced pressure to yield a clear oil. The crude product was dissolved in a minimal amount of DCM, adsorbed onto dry silica, volatiles removed and dry transferred onto a silica column packed in hexanes. The product was eluted with 3 column lengths of 10:90 ethyl acetate: hexanes. The solvent was removed under reduced pressure to yield a clear oil (16 mg, 0.051 mmol, 16% yield). GC-MS *m/z* = 322.39 [M]^+^. ^1^H NMR (500 MHz, CDCl_3_): δ 2.46 (m, 1H), 2.25 (m, 1H), 1.99 (m, 1H), 1.53 (m, 1H), 1.26 (m, 16H), 0.86 (m, 12H). ^13^C NMR (500 MHz, CDCl_3_): δ178.43, 65.86, 41.11, 31.90, 29.33, 27.96, 22.66, 19.69, 14.07.

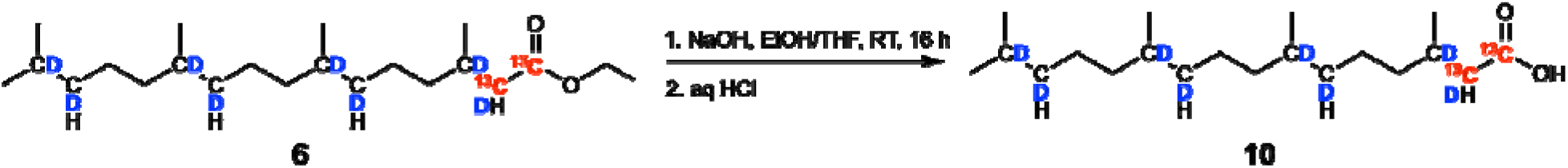

## Results

This is a novel, linear, unified, unoptimized synthesis of all-racemic phytanic acid along with its 2,3,6,7,10,11,14,15-*d*_8_ (**8**); 1,2-^13^*C*_2_ (**9**); and 1,2-^13^*C*_2_-2,3,6,7,10,11,14,15-*d*_8_ (**10**) stable isotopologues from commercially available materials. In the two full schemes below, syntheses of the all-racemic phytanic acid and its three corresponding isotopologues are reported based on the common starting materials, ethyl 2-bromoacetate (**Figure 1**) and ethyl 2-bromoacetate-^13^*C*_2_ (**Figure 2**), respectively. As previously highlighted, phytanic acid may be directly synthesized from natural *trans*-phytol to provide its naturally occurring pair of diastereomers; however, our all-racemic synthesis provided access to the most diverse set of isotopologues from the available starting materials. Although our all-racemic isotopologues of phytanic acid may possess different physicochemical properties from the naturally occurring phytanic acid, the expediency of synthesis by this method was chosen over developing more time-consuming stereoselective synthetic methods or chromatographic separation of stereoisomers. We postulate that cells with functional alpha-, beta-, and omega-oxidation pathways and the alpha-methylacyl-CoA racemase (AMACR) gene may be able to convert the all-racemic phytanic acid to all-racemic pristanic acid, which can subsequently be acted upon by AMACR in the peroxisome.

**Figure 1.**
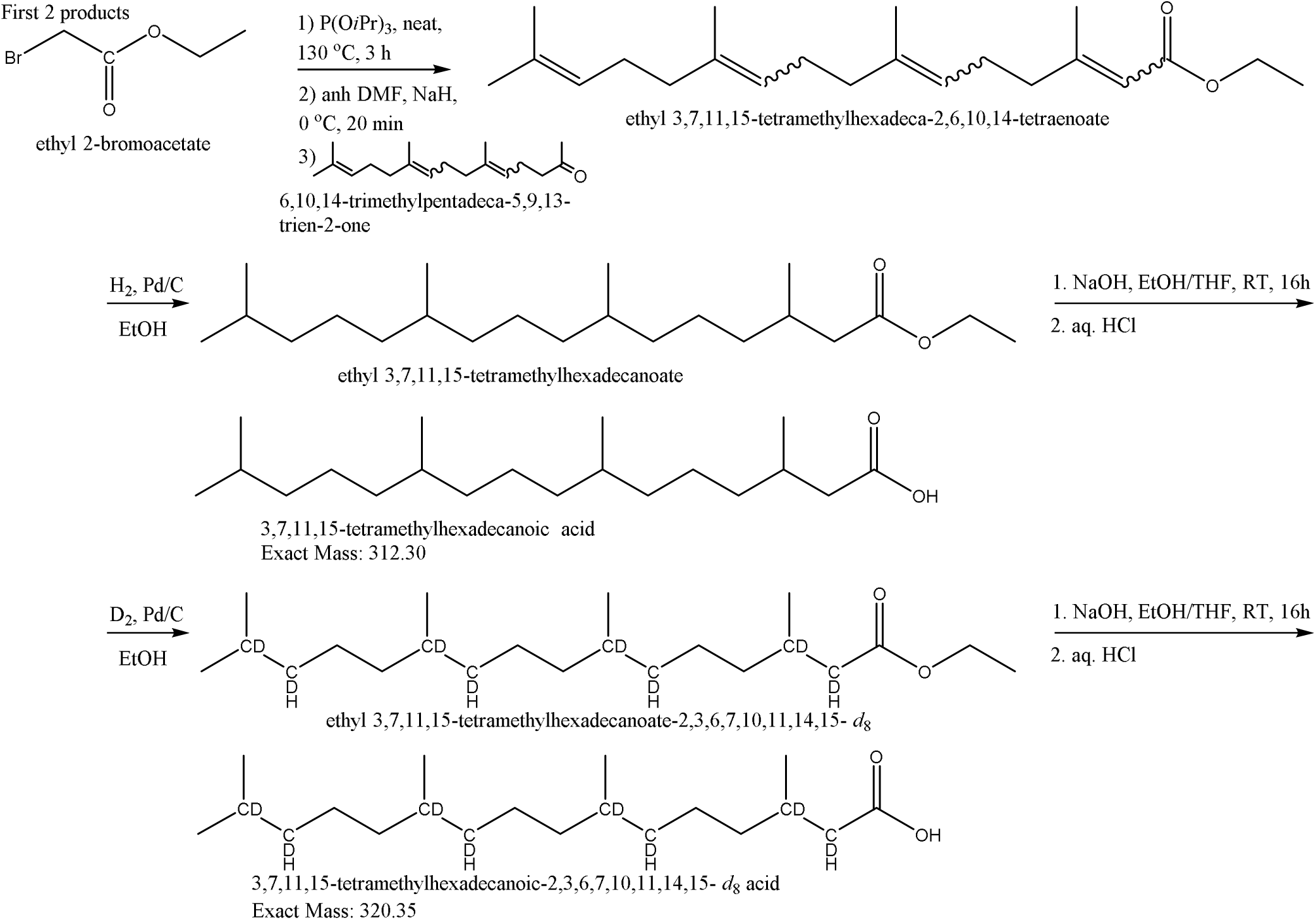
Synthetic route to phytanic acid and 2,3,6,7,10,11,14,15-*d*_8_-phytanic acid.

**Figure 2.**
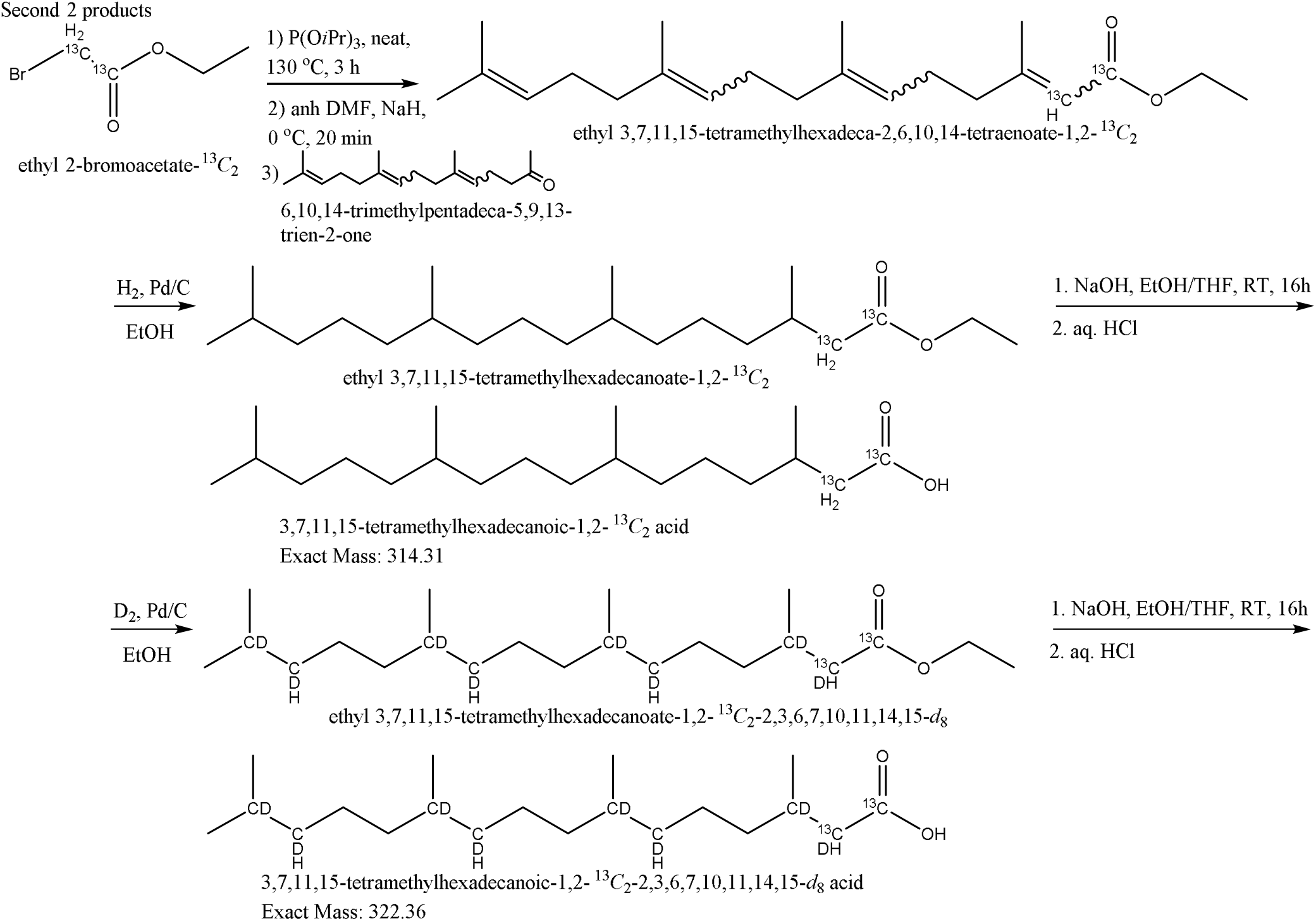
Synthetic route to 1,2-^13^*C*_2_-phytanic acid and 1,2-^13^*C*_2_-2,3,6,7,10,11,14,15-*d*_8_-phytanic acid.

## Acknowledgements/grant support

NWS was supported by R35GM156596 and R01DK138011.

## Author contributions

Conceptualization: RLB; NWS

Synthesis: RLB.; ACS.; JS; DvT

Writing: RLB, NWS

Funding: NWS

## Supporting Information

**Figure S1.**
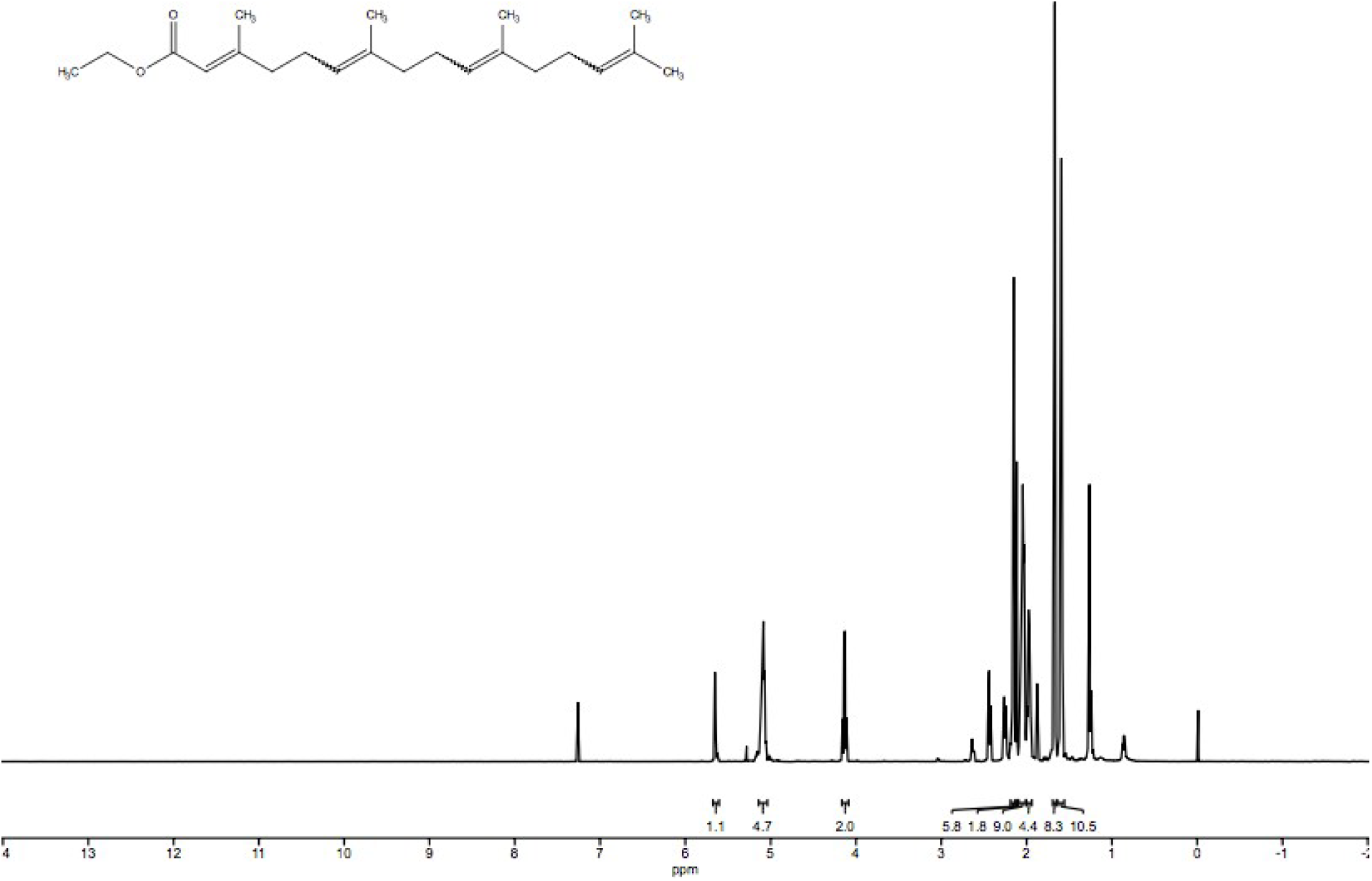
^1^H NMR spectrum for **1**. ^1^H NMR (500 MHz, CDCl_3_): δ 5.65 (s, 1H), 5.09 (m, 4H), 4.14 (m, 2H), 2.15 (m, 6H), 2.12 (s, 2H), 2.04 (m, 9H), 1.98, (m, 4H), 1.67, (m, 8H), 1.59 (t, 10H).

**Figure S2.**
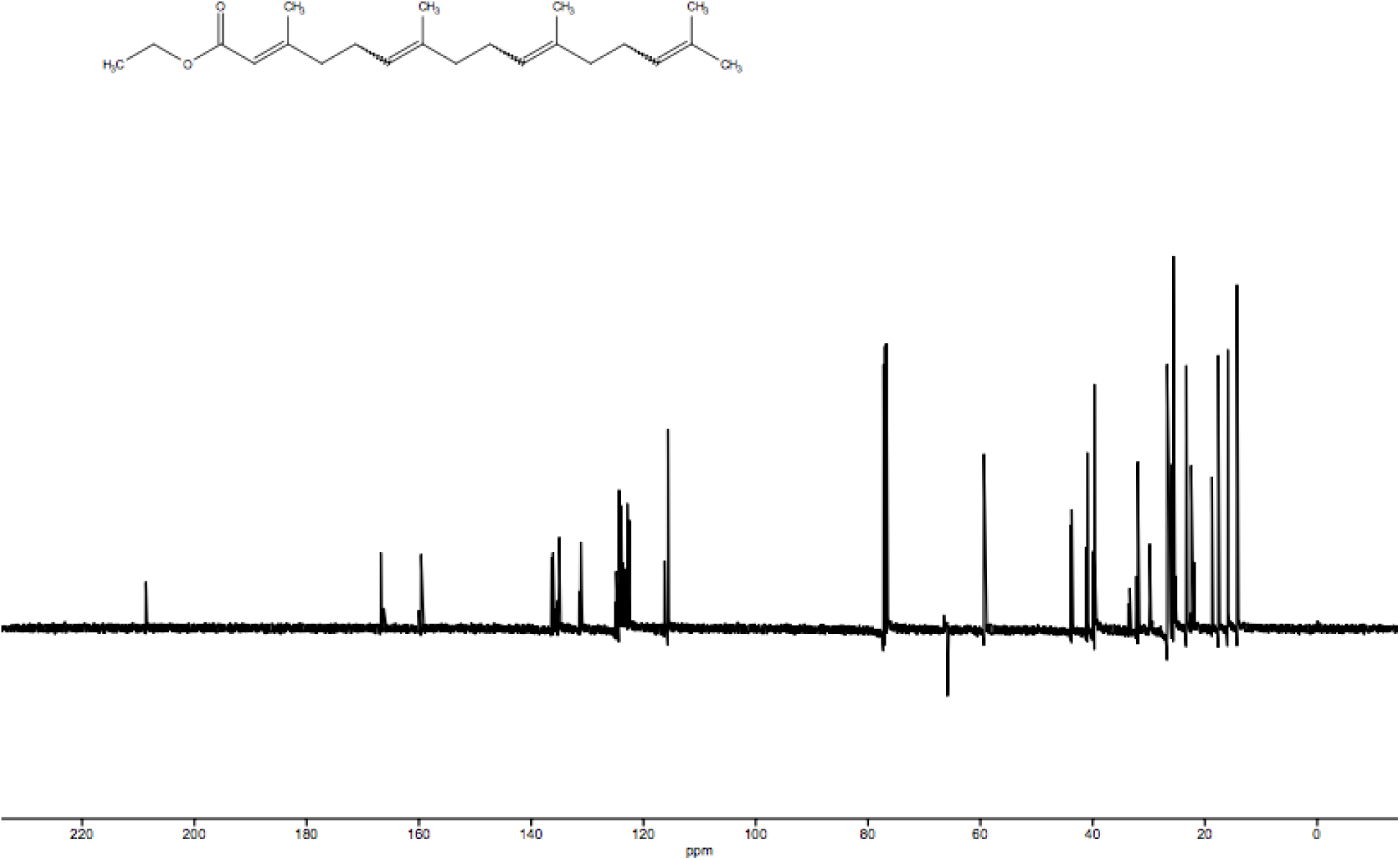
^13^C NMR spectrum for **1**. ^13^C NMR (126 MHz, CDCl_3_): δ 208.63, 166.80, 159.62, 136.35, 134.98, 124.35, 122.89, 115.64, 59.37, 43.73, 40.92, 39.68, 31.96, 26.74, 23.32, 17.62, 14.29.

**Figure S3.**
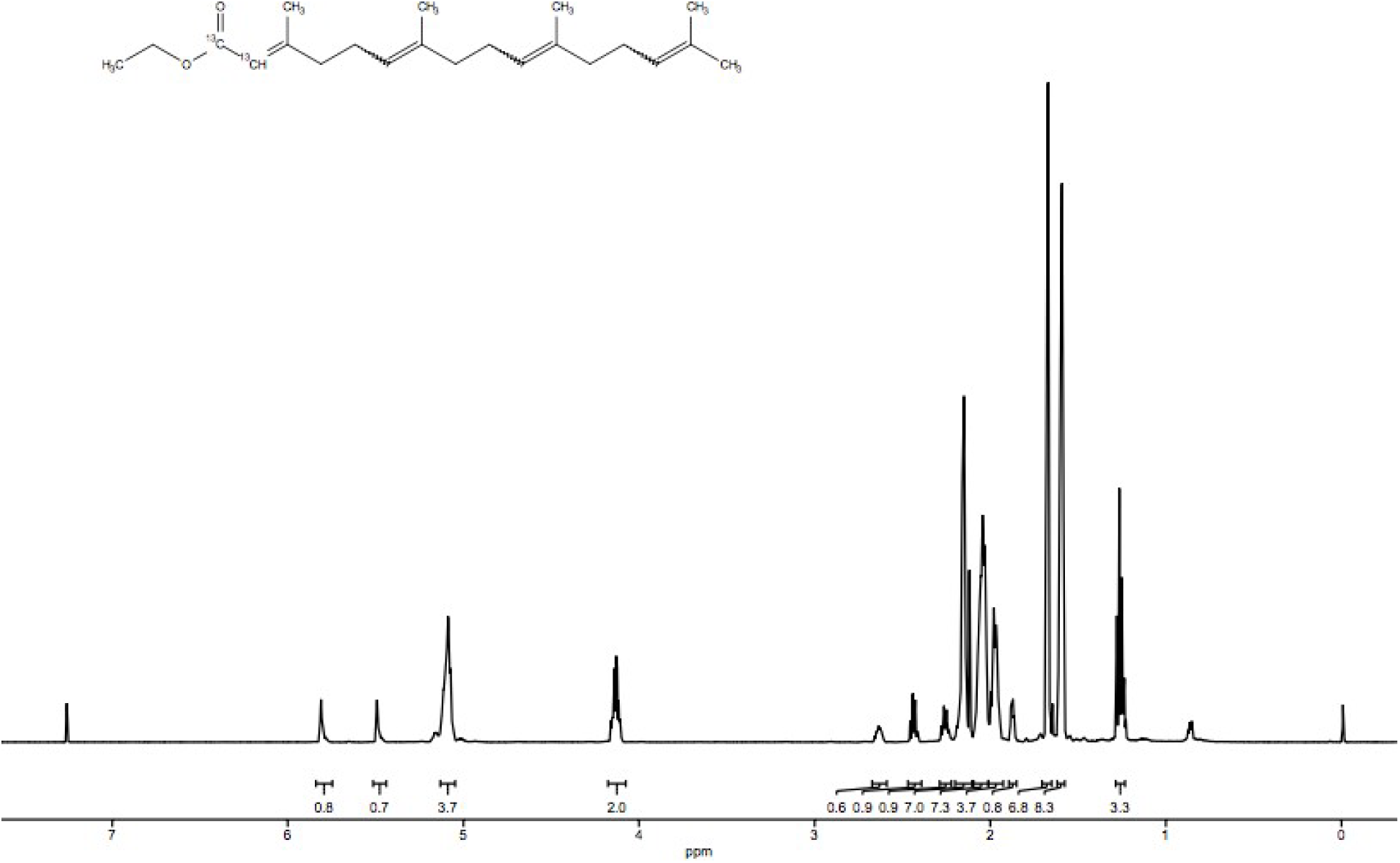
^1^H NMR spectrum for **2**. ^1^H NMR (500 MHz, CDCl_3_): δ 5.81 (s, 1H), 5.49 (s, 1H), 5.09 (t, 4H), 4.12 (m, 2H), 2.65 (m, 1H), 2.44 (m, 1H), 2.26 (q, 1H), 2.15 (s, 7H), 2.04 (m, 7H), 1.97 (m, 4H), 1.67 (s, 7H), 1.59 (s, 8H), 1.24 (m, 3H).

**Figure S4.**
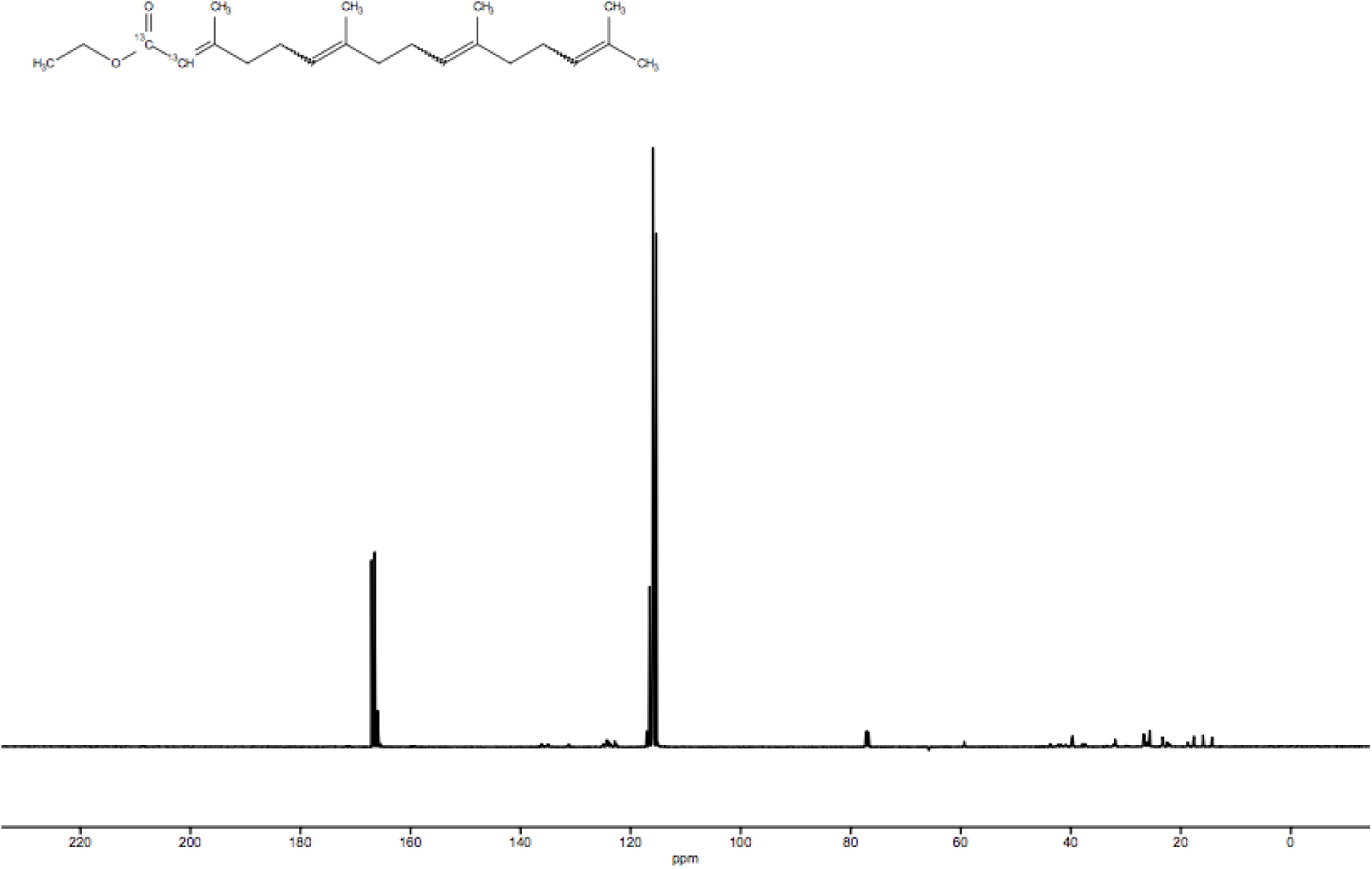
^13^C NMR spectrum for **2**. ^13^C NMR (126 MHz, CDCl_3_): δ167.10, 136.11, 124.09, 115.94, 59.37, 43.74, 41.85, 39.71, 37.85, 31.96, 26.74, 23.32, 17.62, 14.28.

**Figure S5.**
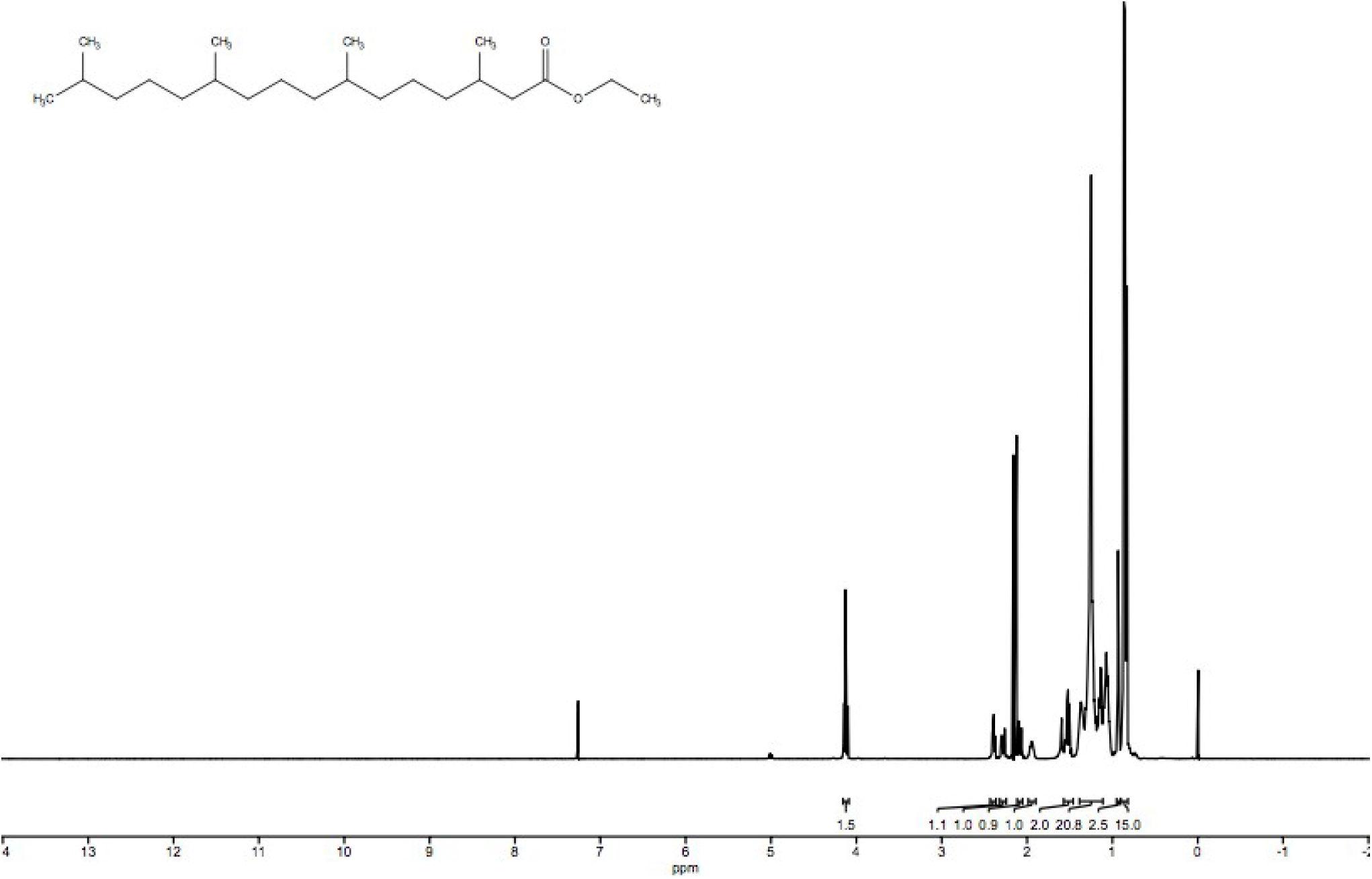
^1^H NMR spectrum for **3.** ^1^H NMR (500 MHz, CDCl_3_): δ 4.11 (q, 2H), 2.39 (t, 1H), 2.27 (m, 1H), 2.07 (m, 1H), 1.95 (m, 1H), 1.52 (m, 2H), 1.25 (m, 20H), 0.93 (m, 2H), 0.86 (m, 15H).

**Figure S6.**
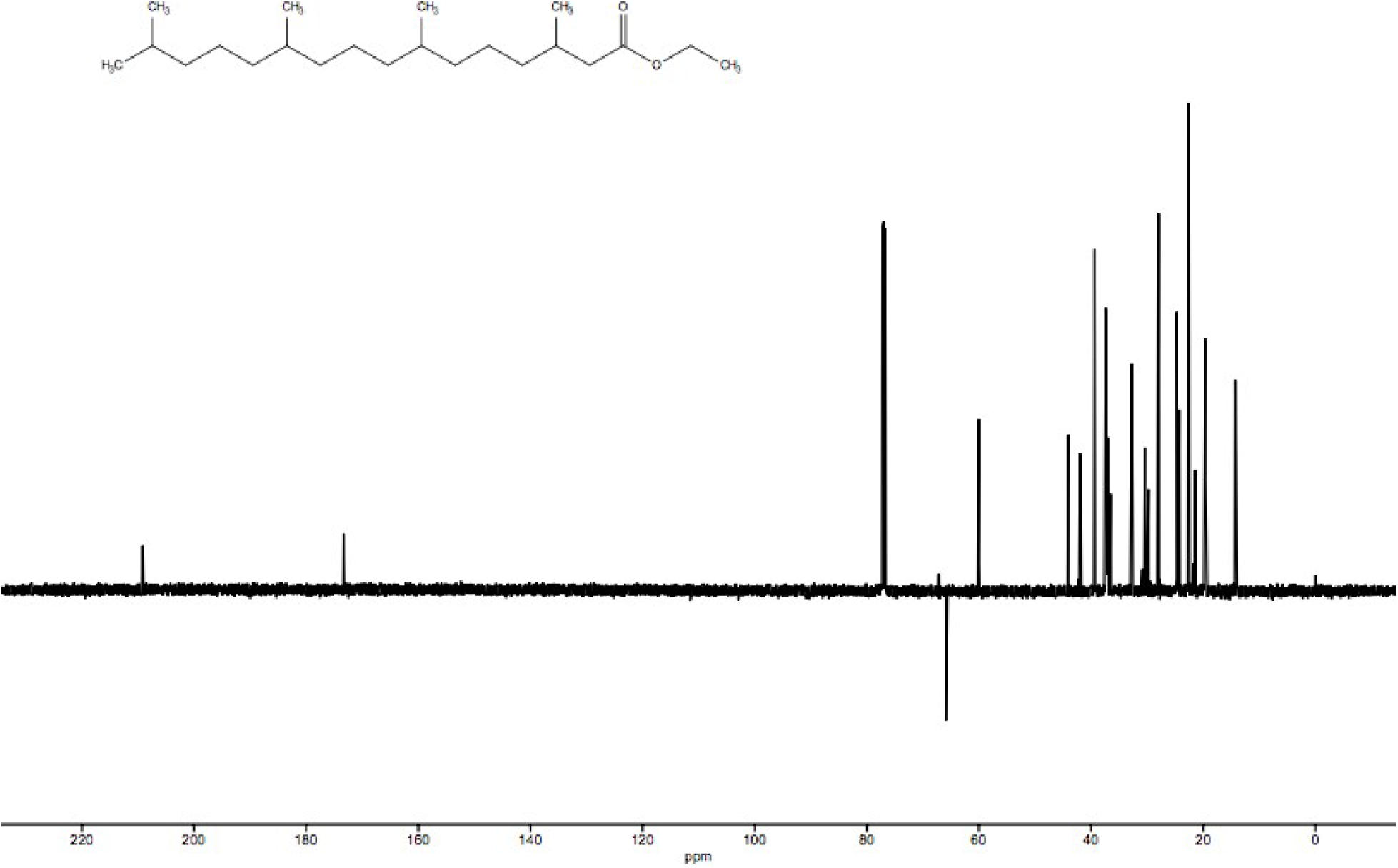
^13^C NMR spectrum for **3.** ^13^C NMR (126 MHz, CDCl_3_): δ 209.18, 173.30, 59.99, 44.11, 41.90, 39.35, 37.36, 32.74, 30.39, 27.95, 24.77, 22.67, 21.44, 19.64, 14.25.

**Figure S7.**
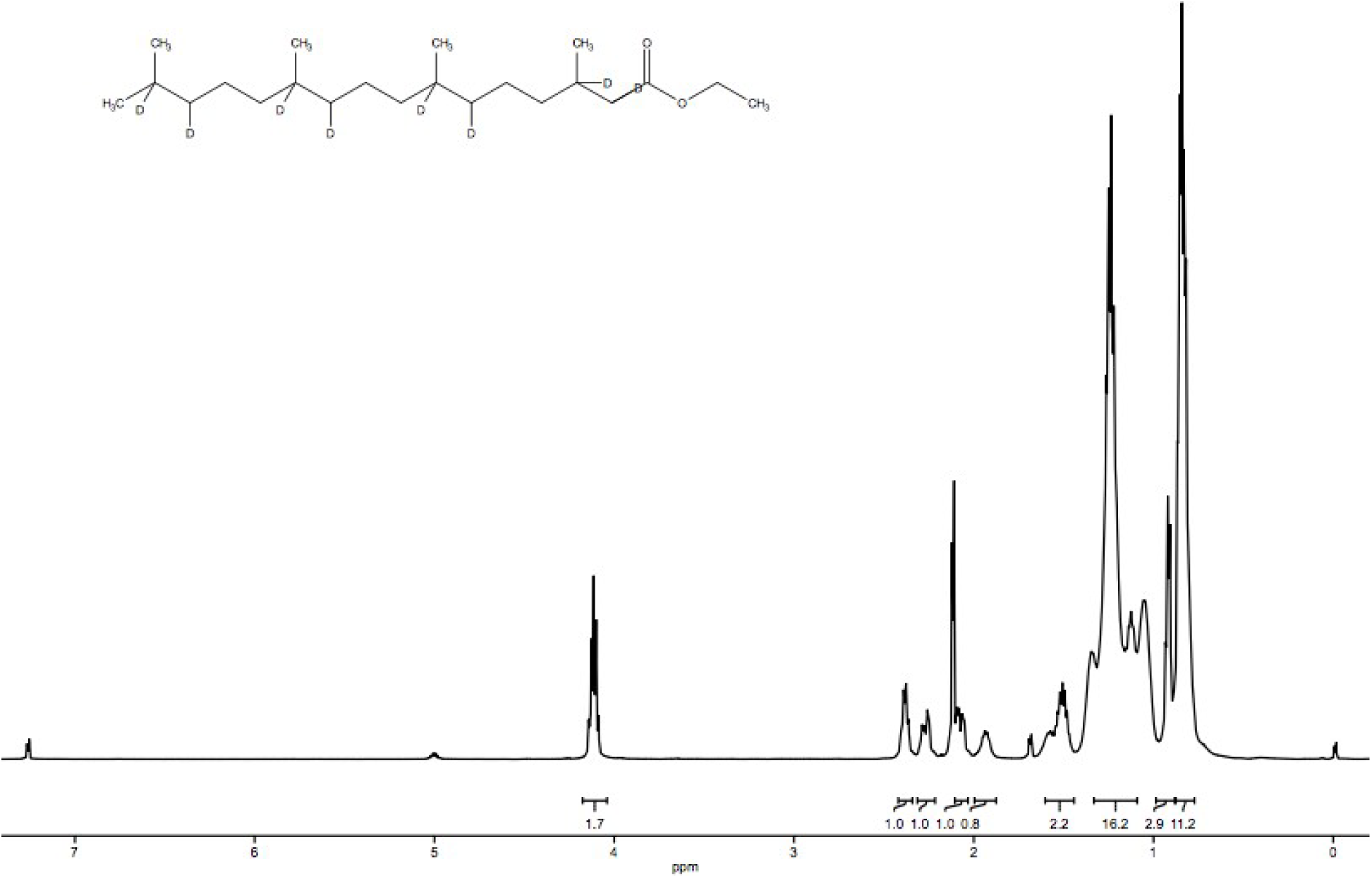
^1^H NMR spectrum for **4**. ^1^H NMR (500 MHz, CDCl_3_): δ 4.11 (q, 2H), 2.39 (q, 1H), 2.29 (m, 1H), 2.05 (m, 1H), 1.91 (m, 1H), 1.52 (m, 2H), 1.25 (m, 16H), 0.92 (t, 3H) 0.84 (m, 11H).

**Figure S8.**
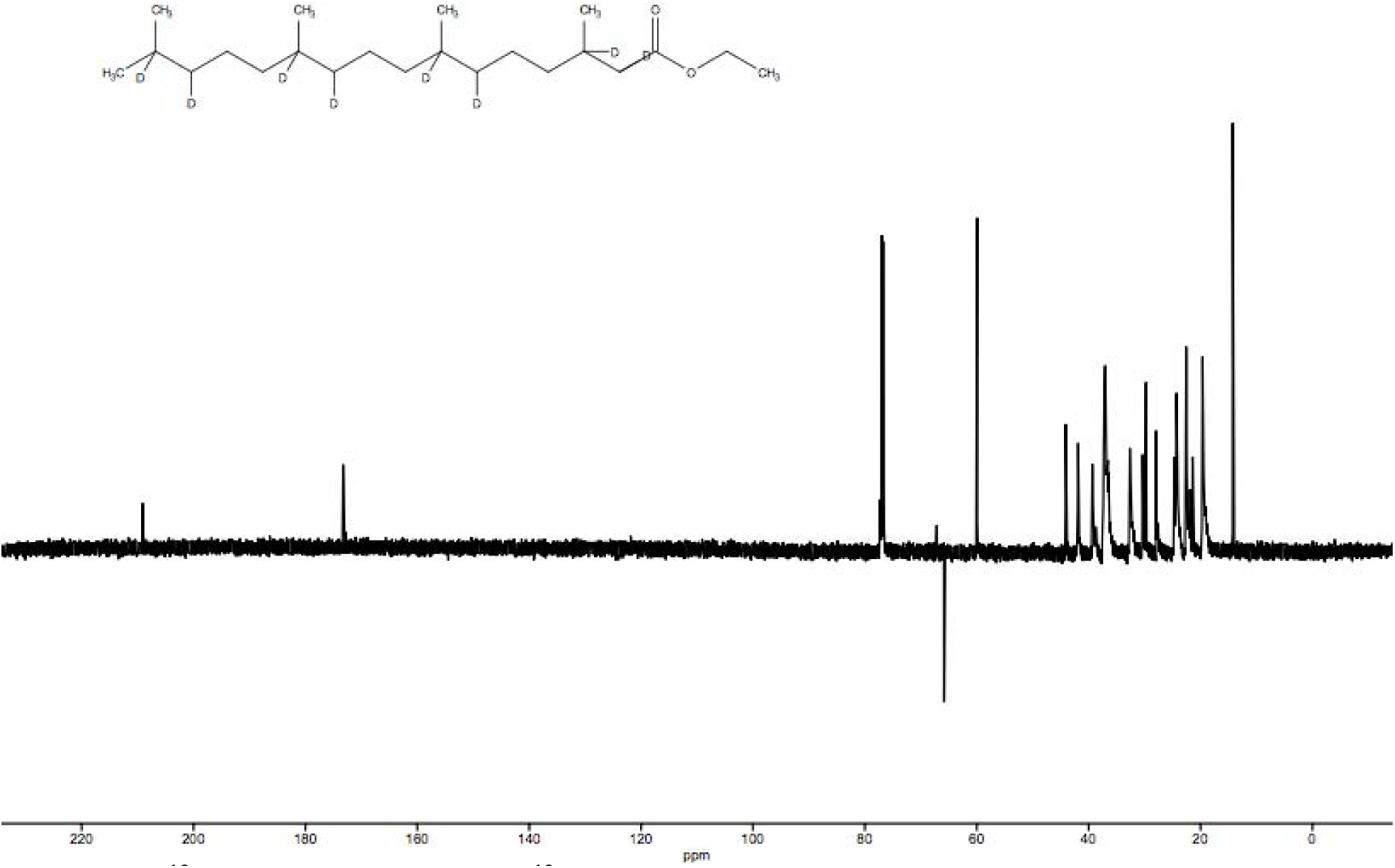
^13^C NMR spectrum for **4**. ^13^C NMR (126 MHz, CDCl_3_): δ 209.09, 173.28, 59.96, 44.09, 41.88, 39.35, 37.04, 29.74, 27.93, 24.28, 22.57, 21.42, 19.61, 14.24.

**Figure S9.**
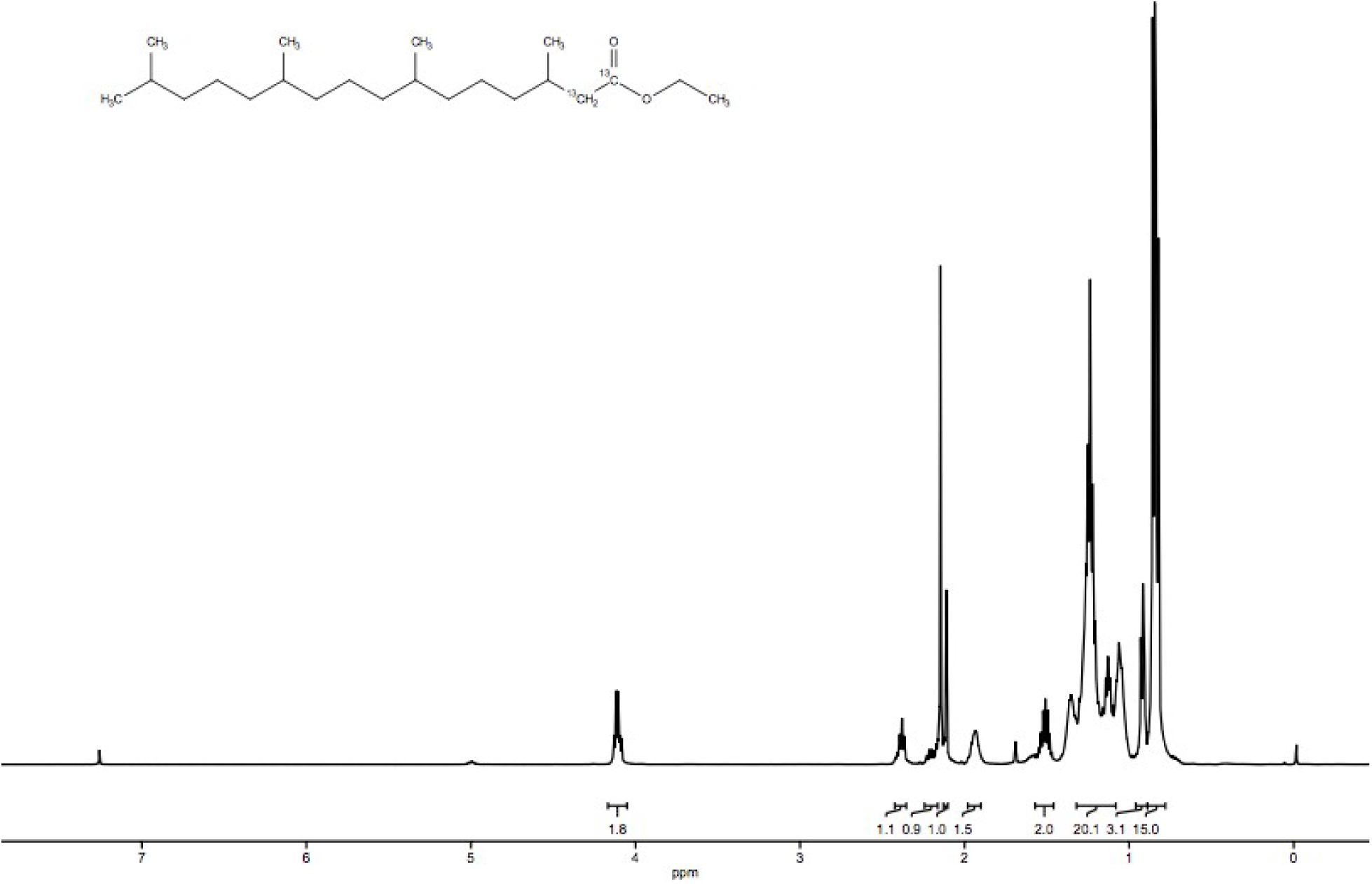
^1^H NMR spectrum for **5**. ^1^H NMR (500 MHz, CDCl_3_): δ 4.10 (q, 2H), 2.38 (t, 1H), 2.15 (m, 1H), 2.11 (s, 1H), 1.93 (m, 1H), 1.51 (m, 2H), 1.24 (m, 20H), 0.92 (t, 3H), 0.84 (m, 15H).

**Figure S10.**
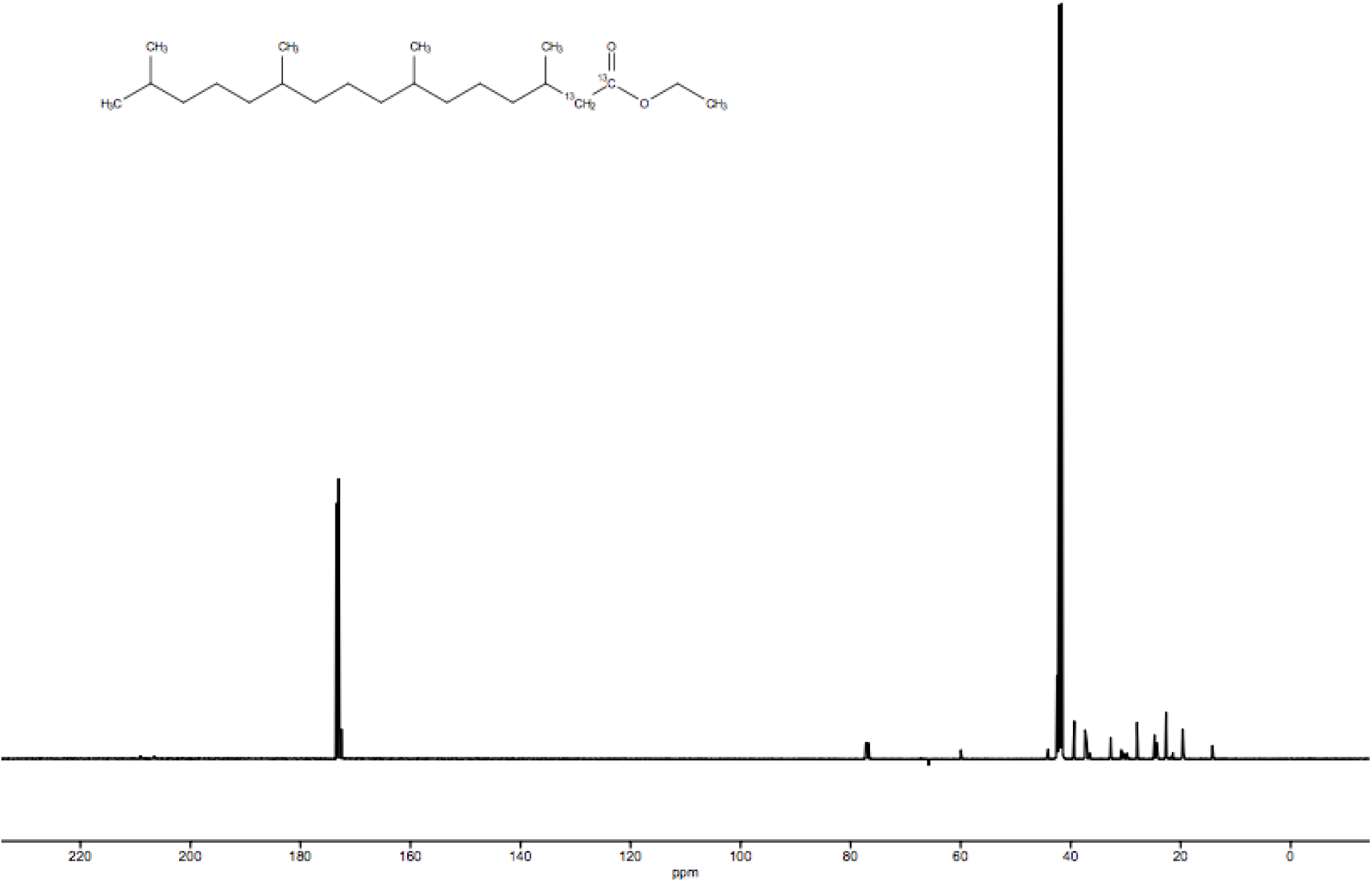
^13^C NMR spectrum for **5**. ^13^C NMR (500 MHz, CDCl_3_): δ 173.03, 59.96, 42.49, 39.34, 37.07, 32.75, 27.93, 24.75, 22.57, 19.62, 14.22.

**Figure S11.**
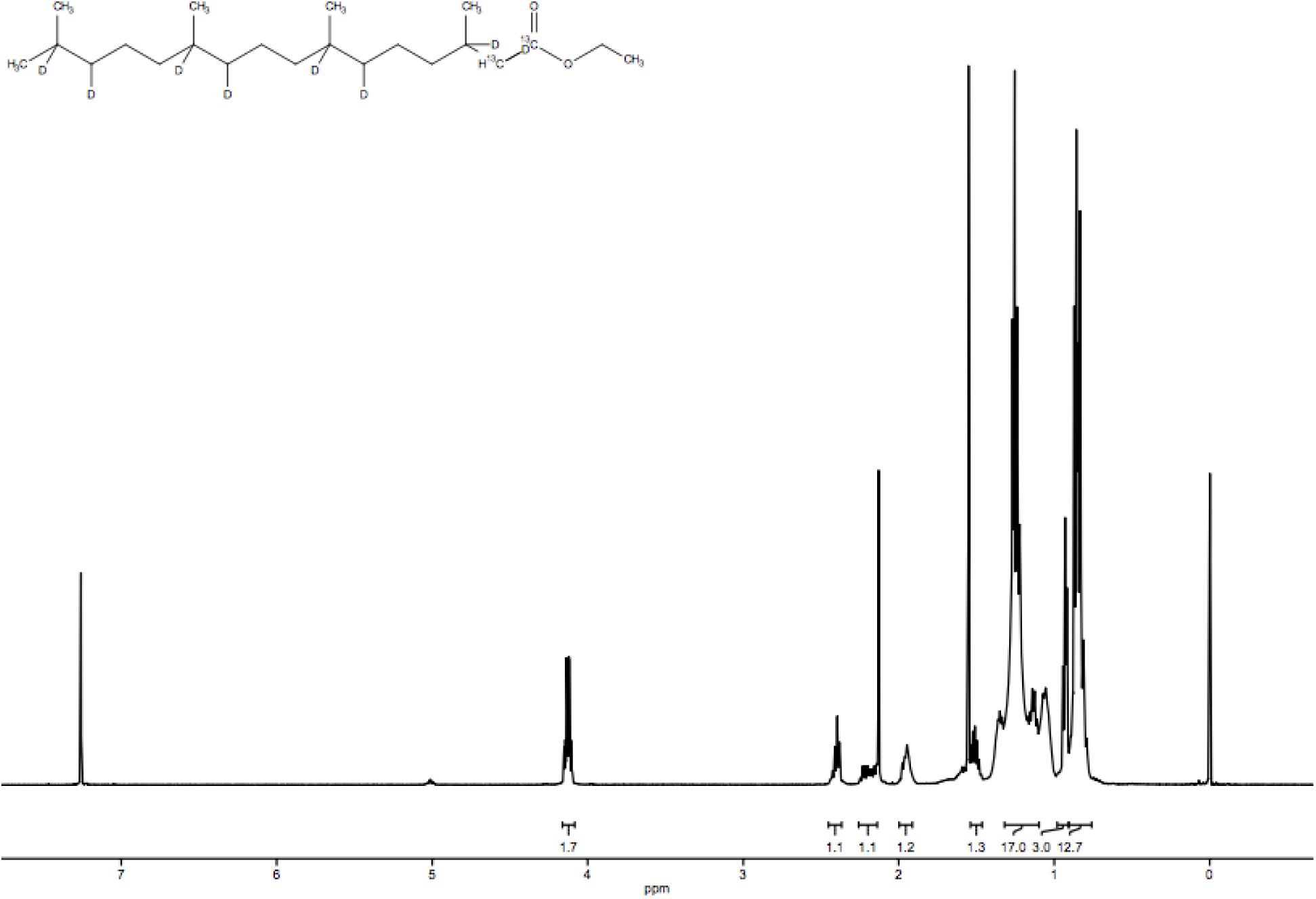
^1^H NMR spectrum for **6**. ^1^H NMR (500 MHz, CDCl_3_): δ 4.12 (m, 2H), 2.39 (t, 1H), 2.20 (m, 1H), 1.95 (m, 1H), 1.51 (m, 1H), 1.25 (m, 17H), 0.93 (t, 3H), 0.86 (m, 13H).

**Figure S12.**
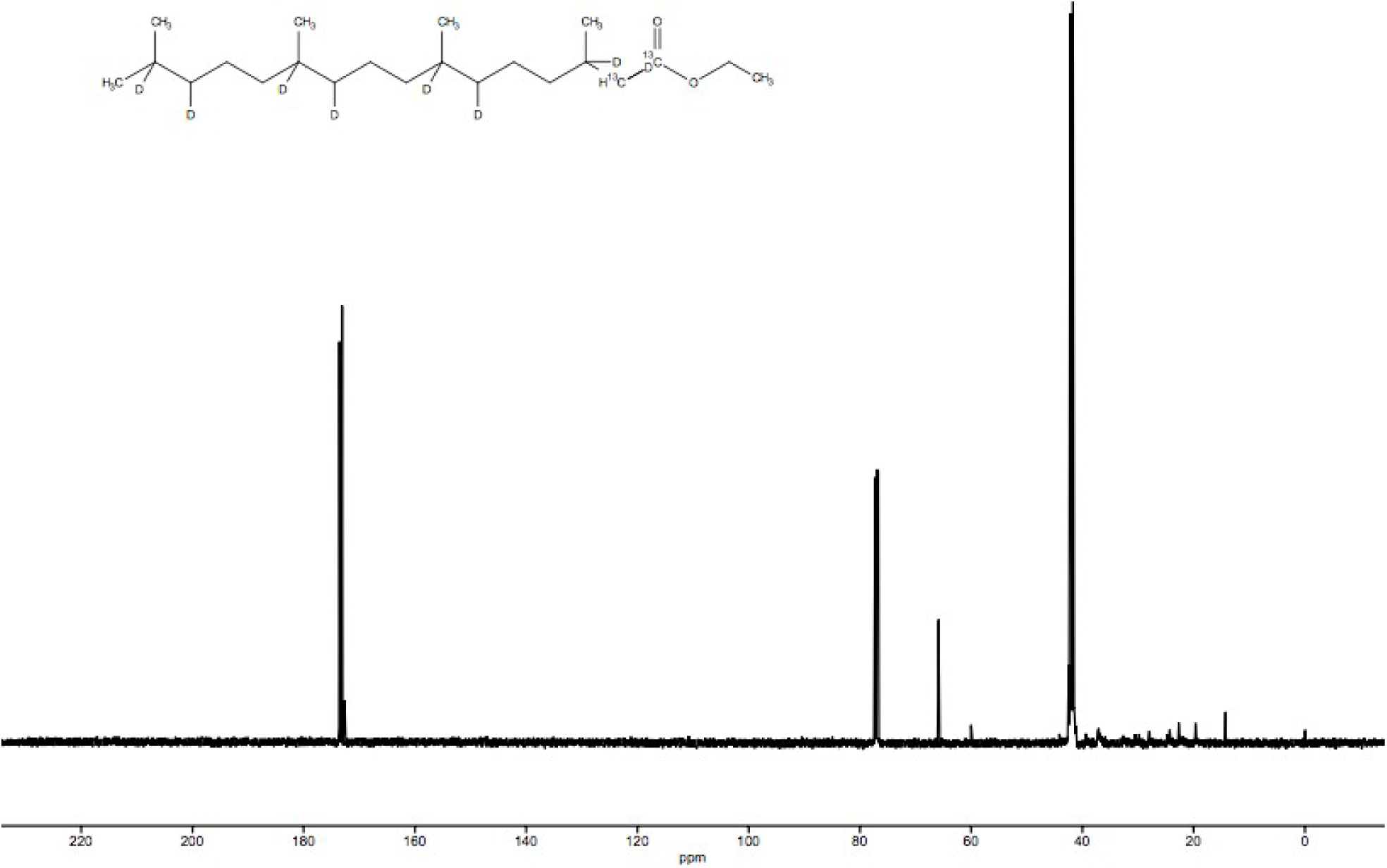
^13^C NMR spectrum for **6**. ^13^C NMR (500 MHz, CDCl_3_): δ 173.12, 65.86, 60.02, 41.68, 27.96, 22.68, 19.66, 14.27.

**Figure S13.**
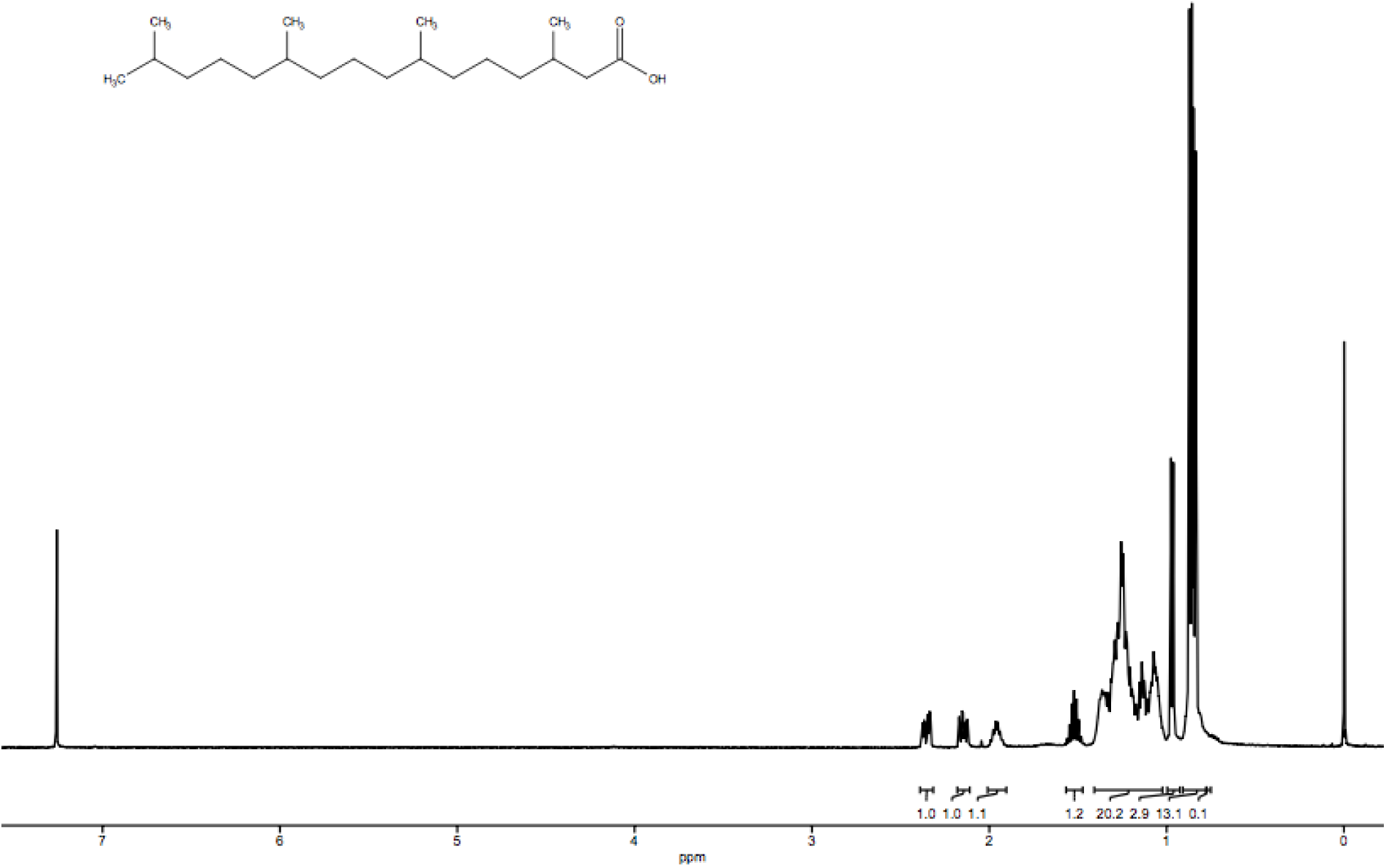
^1^H NMR spectrum for **7**. ^1^H NMR (500 MHz, CDCl_3_): δ 2.35 (m,1H), 2.12 (m, 1H), 1.95 (m, 1H), 1.52 (m, 1H), 1.26 (m, 20H), 0.96 (d, 3H), 0.87, (q, 13H). (Dora von Trentini)

**Figure S14.**
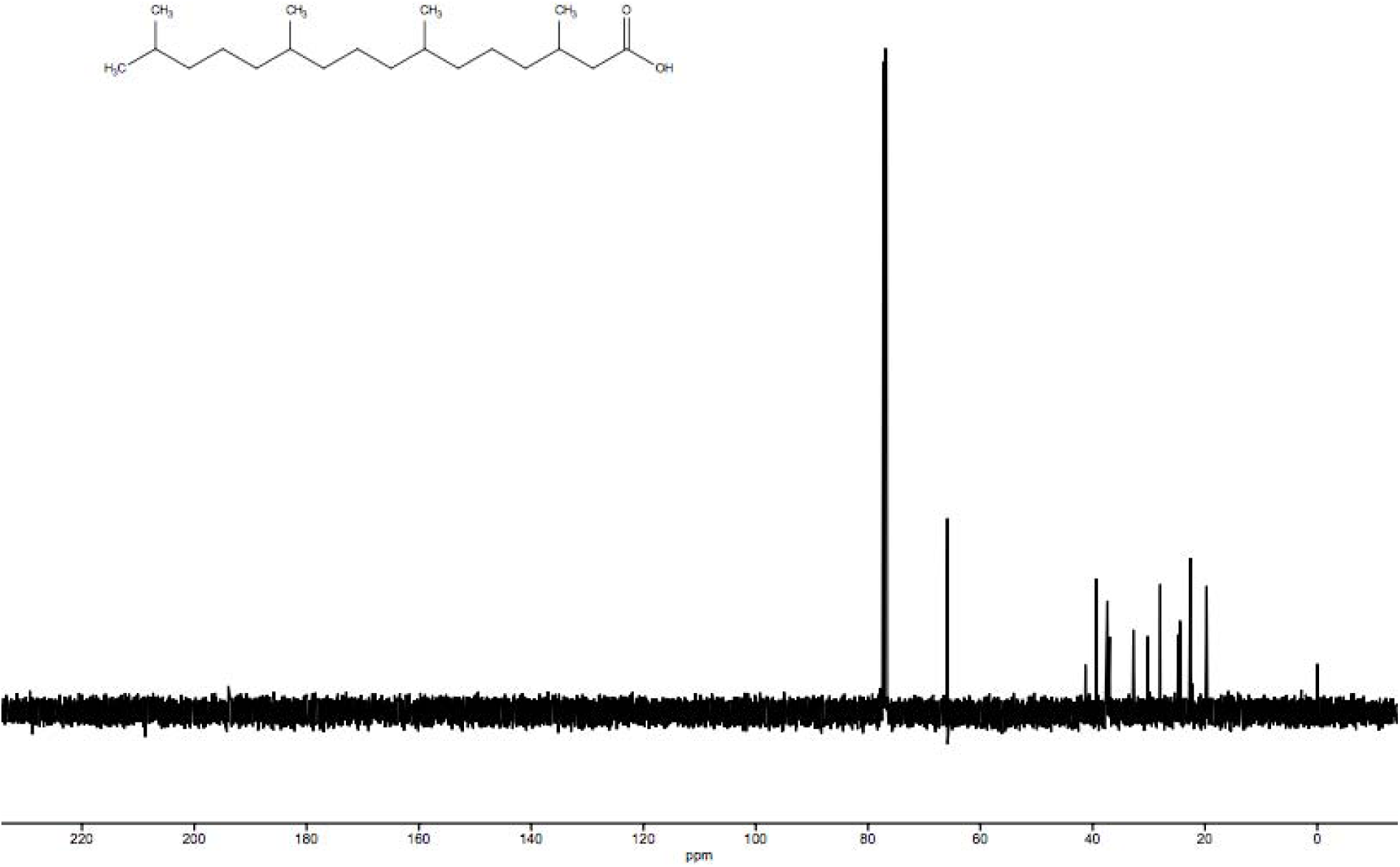
^13^C NMR spectrum for **7**. ^13^C NMR (500 MHz, CDCl_3_): δ 65.86, 41.38, 39.36, 37.27, 32.76, 30.18, 27.97, 24.32, 22.62, 19.64.

**Figure S15.**
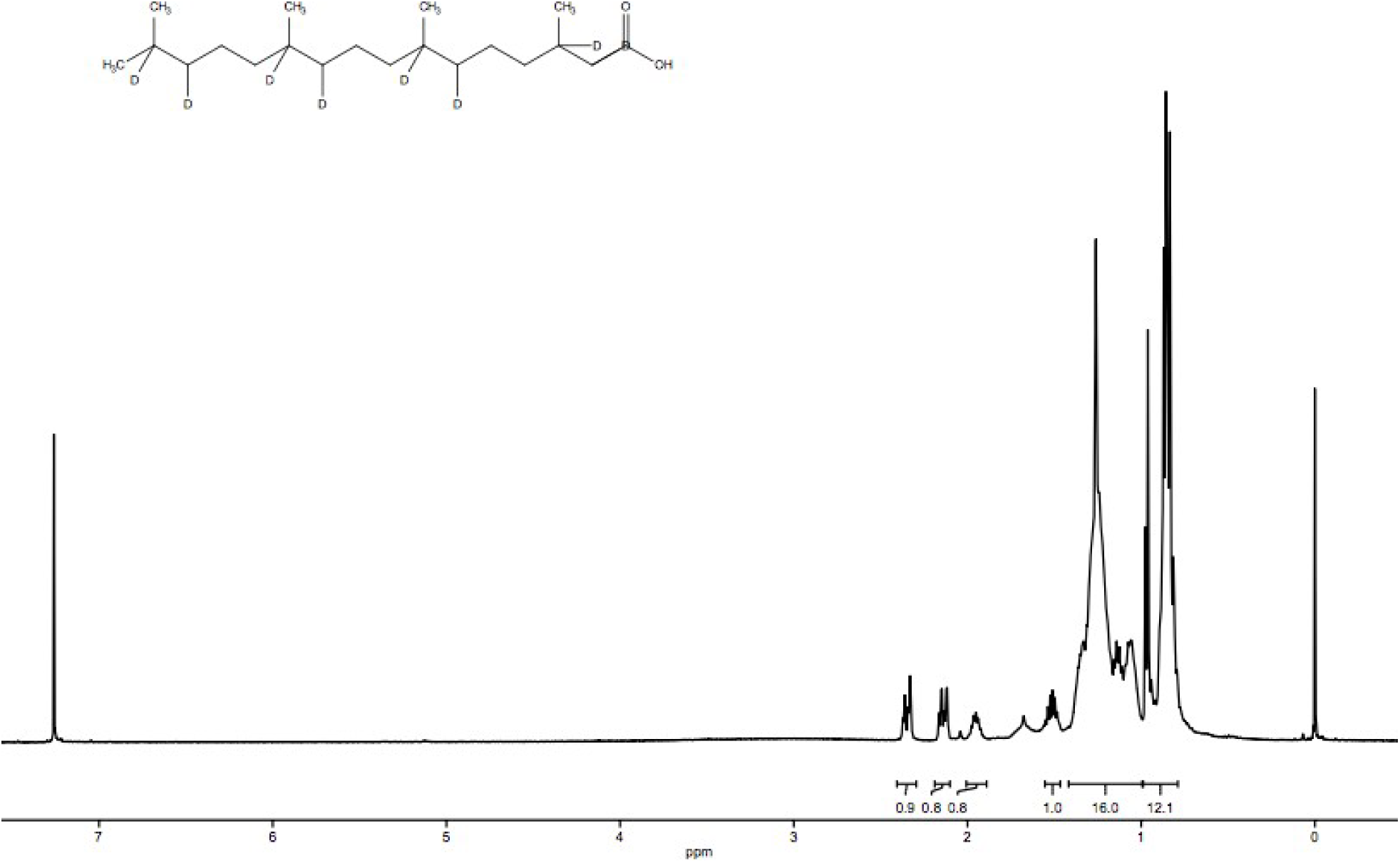
^1^H NMR spectrum for **8**. ^1^H NMR (500 MHz, CDCl_3_): δ 2.33 (dd, 1H), 2.15 (m, 1H), 1.95 (m, 1H), 1.51 (m, 1H), 1.26 (m, 16H), 0.86 (m, 12 H).

**Figure S16.**
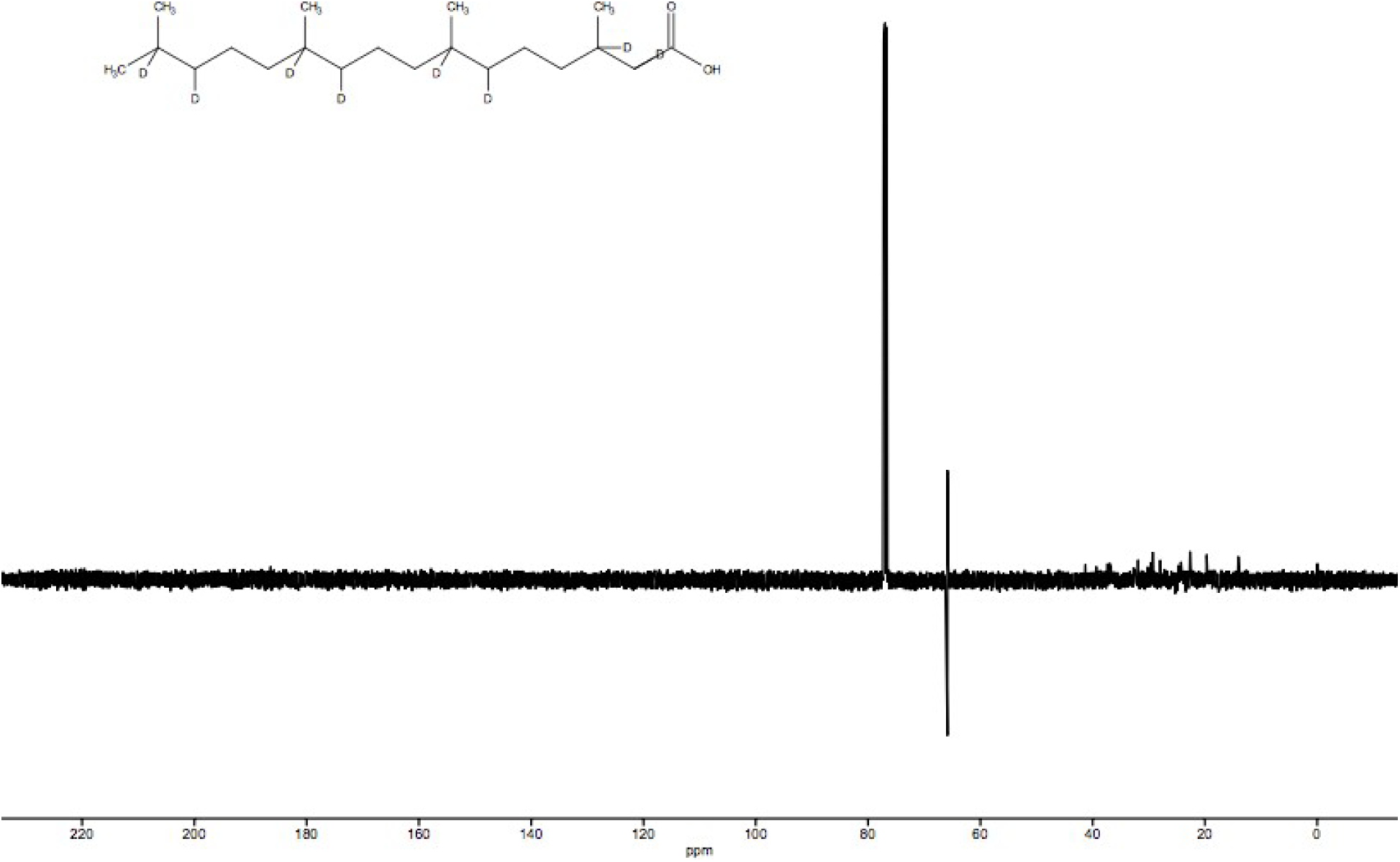
^13^C NMR spectrum for **8.** ^13^C NMR (500 MHz, CDCl_3_): δ 41.29, 37.02, 31.90, 29.33, 24.78, 22.68, 19.51, 14.07.

**Figure S17.**
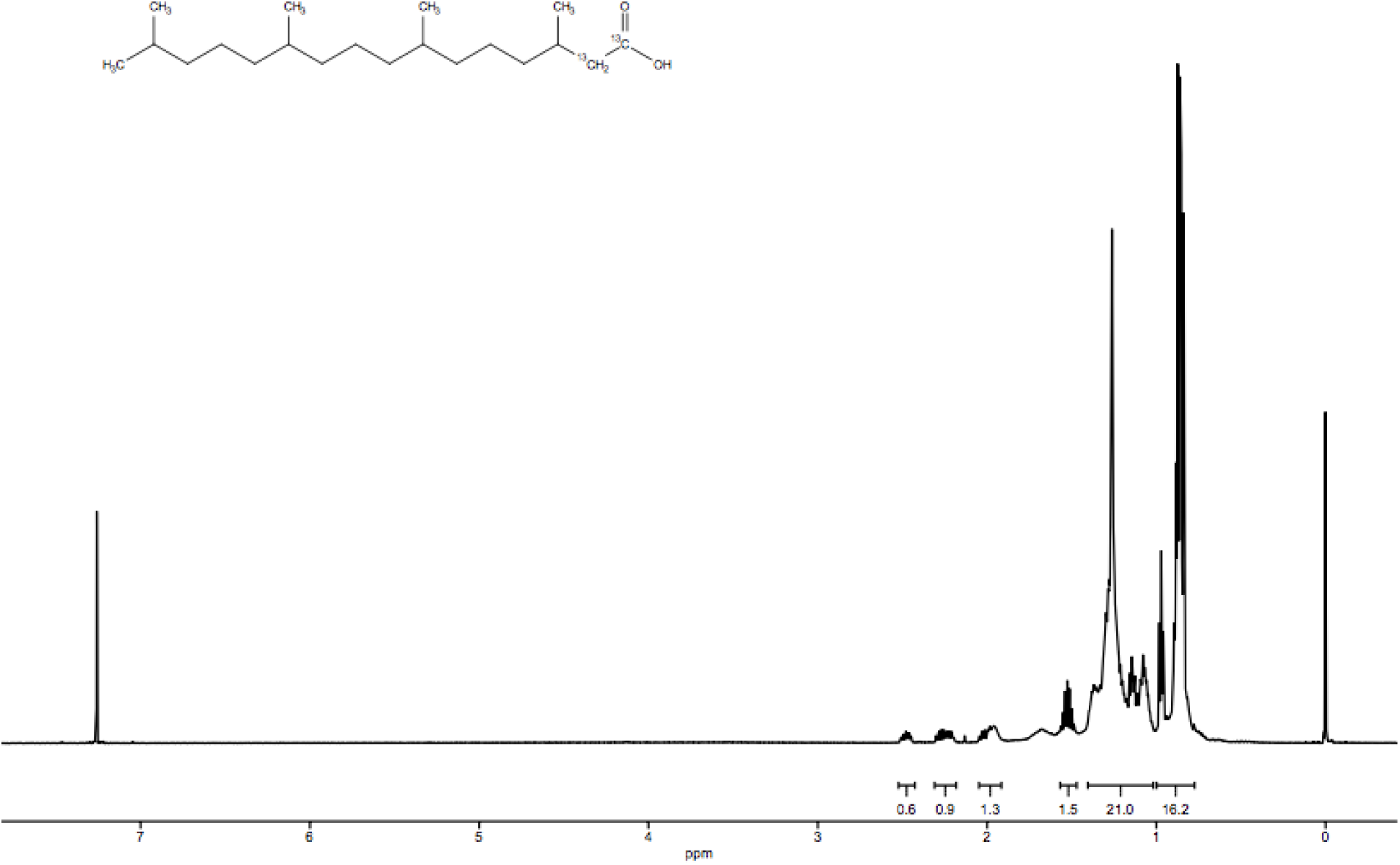
^1^H NMR spectrum for **9**. ^1^H NMR (500 MHz, CDCl_3_): δ 2.48 (m, 1H), 2.25 (m, 1H), 2.00, (m, 1H), 1.53 (m, 1H), 1.26 (m, 21H), 0.86 (m, 16H).

**Figure S18.**
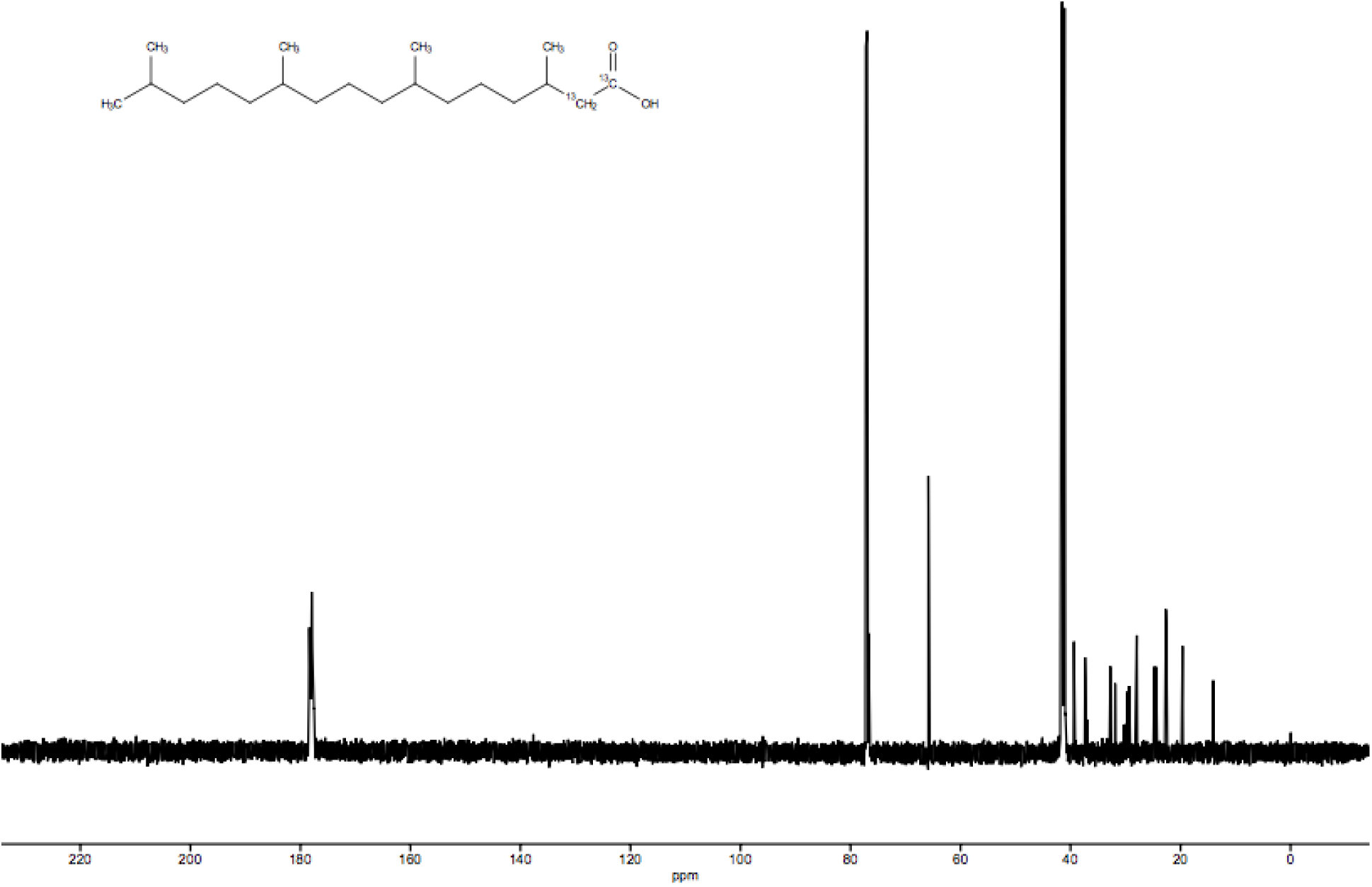
^13^C NMR spectrum for **9**. ^13^C NMR (500 MHz, CDCl_3_): δ 177.90, 65.86, 4108, 39.36, 37.37, 32.78, 31.90, 29.33, 27.96, 24.77, 22.68, 19.64, 14.07.

**Figure S19.**
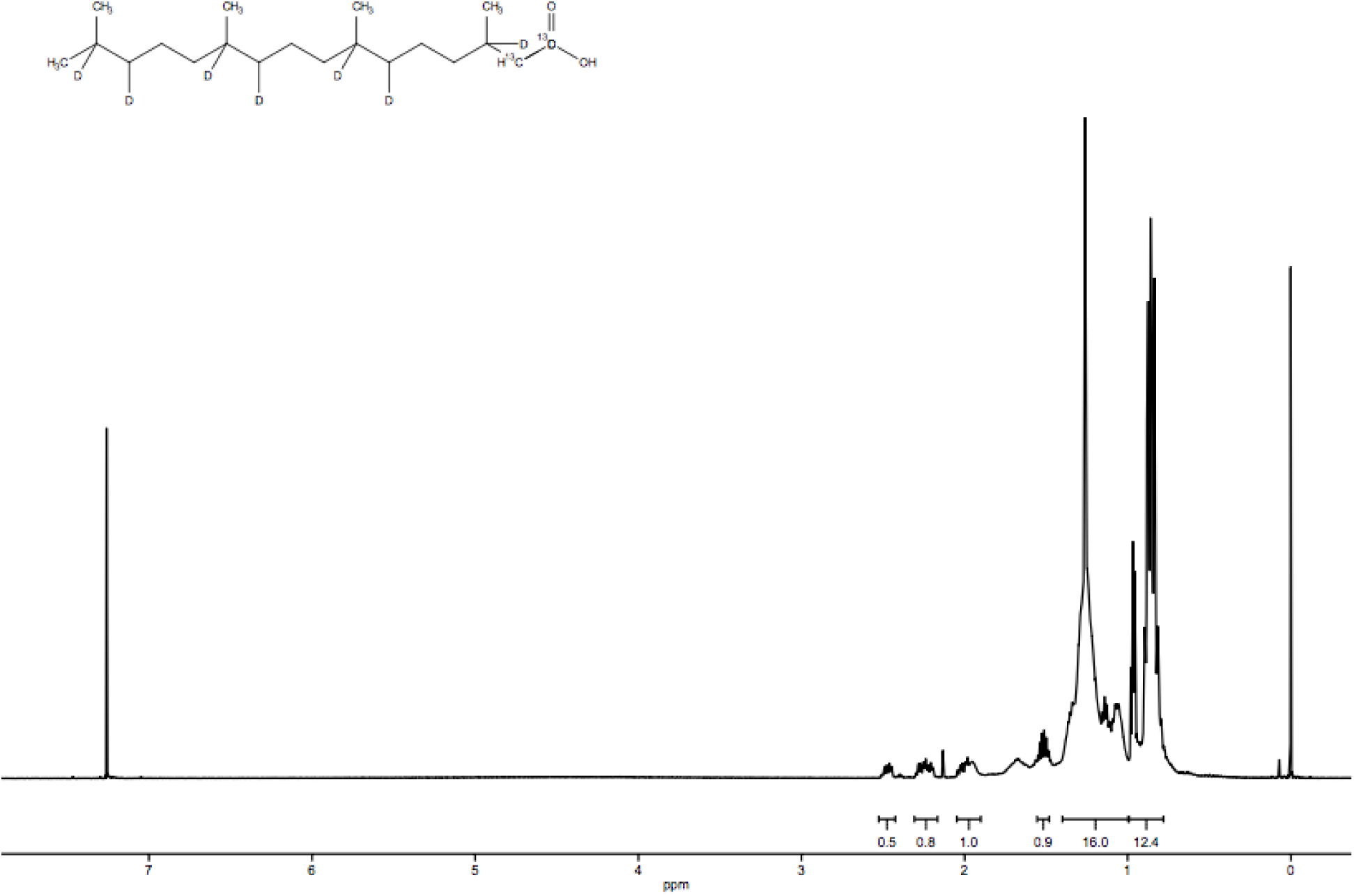
^1^H NMR spectrum for **10.** ^1^H NMR (500 MHz, CDCl_3_): δ 2.46 (m, 1H), 2.25 (m, 1H), 1.99 (m, 1H), 1.53 (m, 1H), 1.26 (m, 16H), 0.86 (m, 12H).

**Figure S20.**
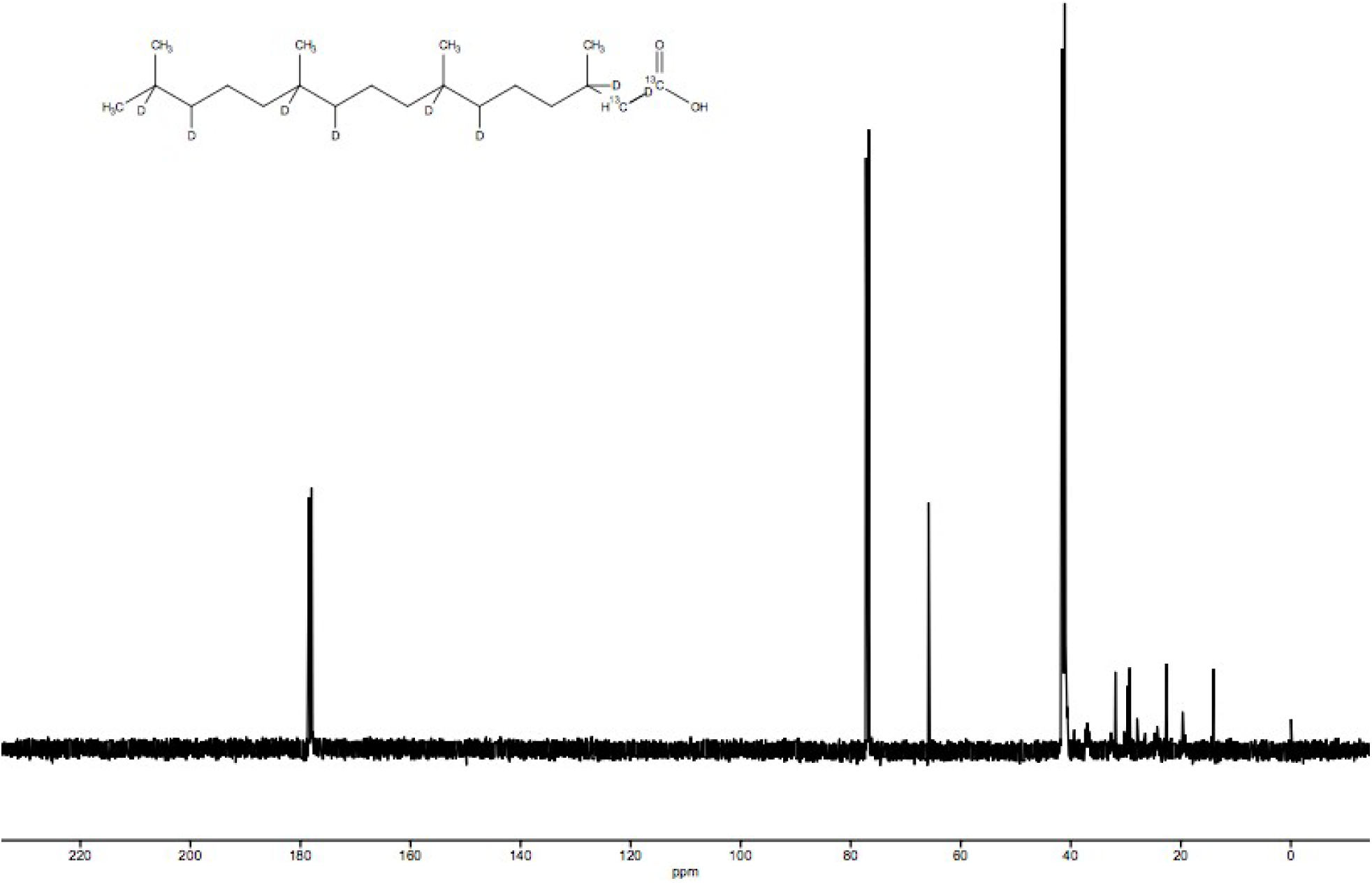
^13^C NMR spectrum for **10.** ^13^C NMR (500 MHz, CDCl_3_): δ 178.43, 65.86, 41.11, 31.90, 29.33, 27.96, 22.66, 19.69, 14.07.

**Figure S21.**
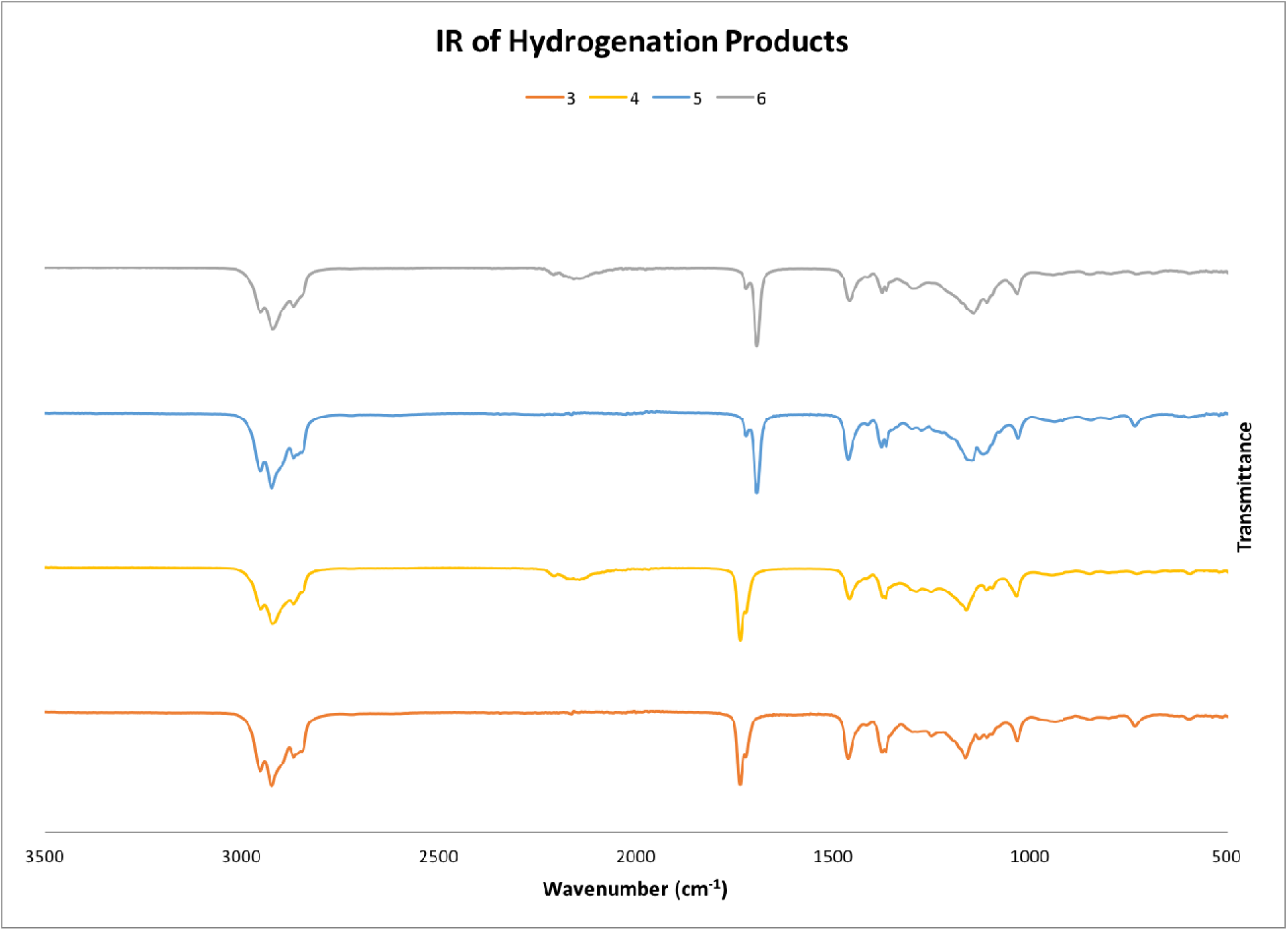
IR spectra of **3, 4, 5**, and **6**.

